# Directed evolution-based discovery of ligands for in vivo restimulation of CAR-T cells

**DOI:** 10.1101/2024.04.16.589780

**Authors:** Tomasz M. Grzywa, Alexandra Neeser, Ranjani Ramasubramanian, Anna Romanov, Ryan Tannir, Naveen K. Mehta, Benjamin Cossette, Duncan M. Morgan, Beatriz Goncalves, Ina Sukaj, Elisa Bergaggio, Stephan Kadauke, Regina M. Myers, Luca Paruzzo, Guido Ghilardi, Austin Cozzone, Stephen J. Schuster, Noelle Frey, Libin Zhang, Parisa Yousefpour, Wuhbet Abraham, Heikyung Suh, Marco Ruella, Stephan A. Grupp, Roberto Chiarle, K. Dane Wittrup, Leyuan Ma, Darrell J. Irvine

**Author notes:** Correspondence (L.M.), (D.J.I). These authors contributed equally (Alexandra Neeser, Ranjani Ramasubramanian).

## Abstract

Chimeric antigen receptor (CAR) T cell therapy targeting CD19 elicits remarkable clinical efficacy in B-cell malignancies, but many patients relapse due to failed expansion and/or progressive loss of CAR-T cells. We recently reported a strategy to potently restimulate CAR-T cells *in vivo*, enhancing their functionality by administration of a vaccine-like stimulus comprised of surrogate peptide ligands for a CAR linked to a lymph node-targeting amphiphilic PEG-lipid (termed CAR-T-vax). Here, we demonstrate a general strategy to generate and optimize peptide mimotopes enabling CAR-T-vax generation for any CAR. Using the clinical CD19 CAR (FMC63) as a test case, we employed yeast surface display to identify peptide binders to soluble IgG versions of FMC63, which were subsequently affinity matured by directed evolution. CAR-T vaccines using these optimized mimotopes triggered marked expansion and memory development of CD19 CAR-T cells in both syngeneic and humanized mouse models of B-ALL/Lymphoma, and enhanced control of disease progression. This approach thus enables vaccine boosting to be applied to any clinically relevant CAR-T cell product.

## Introduction

CD19-targeted chimeric antigen receptor (CAR) T cell therapy elicits impressive anti-tumor activity in the treatment of relapsed or refractory B-cell acute lymphoblastic leukemia (B-ALL) and lymphoma in both pediatric and adult patients^1,2^. This clinical success led to the FDA approval of four CD19-targeted CAR-T products, namely TECARTUS, KYMRIAH, YESCARTA, and BREYANZI^1^. However, a significant fraction (30-60%) of patients still experience leukemia relapse, among which ∼50% are CD19-positive relapses^3^, suggesting impaired function and/or persistence of the infused CAR-T cells. Robust CAR-T cell engraftment and expansion *in vivo* is a prerequisite for antitumor efficacy^4^, a feature that is affected by several characteristics of the CAR-T cell product such as T cell quality^5^, CAR designs^3,4,6^ and the proportion of early memory T cells in the product^7^. In current CAR-T cell therapy treatment regimens, patients usually receive lymphodepletion to promote CAR-T cell engraftment^8^. Interestingly, multiple reports have shown a correlation between the early-stage expansion of CAR-T cells and initial tumor burden^8,9^. These findings suggest the capacity of CAR-T cells to receive antigen-specific stimulation post-infusion is critical for CAR-T cell expansion, engraftment, and potentially long-term persistence. However, whether the antigens presented by tumor cells are sufficient to optimally stimulate CAR-T cells remains unclear.

Vaccination is a natural process whereby T-cells receive antigen-specific stimulation leading to proliferation, differentiation, and induction of effector functions^10^– effects which could be beneficial in promoting CAR-T cell function *in vivo*. However, traditional vaccines provide antigens that must be processed into peptides by professional antigen-presenting cells (APCs) and presented on the major histocompatibility complex (MHC) molecules to stimulate T-cells through the T-cell receptor (TCR). This is challenging to adapt to the setting of engineered CAR-T cells, as current CAR-T products are polyclonal and express uncharacterized TCRs. Further, recent next-generation engineering efforts such as knocking CARs into the TCR locus produce CAR-T lacking endogenous TCRs^11^. Proof of concept demonstration of CAR-T cells generated from virus-specific T cells followed by vaccine boosting via the viral-specific TCR has been shown in small clinical trials, yet the benefits observed with this approach have been limited^12,13^.

To overcome these issues, we recently developed a molecularly-targeted amphiphilic vaccine (amph-vax) approach to directly stimulate CAR-T cells through the chimeric antigen receptor^14,15^. Amph-vax molecules were generated by linking a CAR-ligand to albumin-binding phospholipid polymers. Following parenteral injection, these CAR-ligand conjugates associate with endogenous albumin present in interstitial fluid and are carried into lymph and downstream draining lymph nodes (dLNs). In the dLN, the lipid tails of amph-vax ligands insert into cell membranes, including membranes of professional APCs, allowing APCs to directly “present” the CAR ligand to CAR-T cells together with additional co-stimulation and cytokine support, making them *de novo* CAR-T cell-priming APCs (**Extended Data Fig. 1**)^14^. Amph-vax stimulation led to pronounced CAR-T cell expansion with enhanced functionality, memory formation and substantially enhanced tumor clearance^14,15^. Importantly, we also found that vaccine boosting of CAR-T cells promotes antigen spreading and the induction of endogenous anti-tumor immune responses, which prevent antigen loss-mediated tumor escape^15^. We demonstrated these effects using CARs that recognize synthetic ligands (e.g., fluorescein) and a model EGFRvIII-specific CAR, where the scFv domain of the chimeric receptor recognized a linear peptide epitope in EGFRvIII that is readily synthesized as an amph-ligand. However, to generalize this strategy and enable amph-vax boosting with any arbitrary CAR of interest, an approach is needed for *de novo* generation of surrogate peptide ligands for chimeric antigen receptors.

Here, we demonstrate a workflow to identify peptide mimotope ligands for CAR receptors using yeast surface display-based directed evolution^16^, which allows for the generation of amph-vaccines that can be used with any CAR system. We demonstrate the pipeline by discovering an amph-ligand that can be used with all four FDA-approved CD19 CAR-T cell therapies. We created a customized mimotope yeast library to identify peptides that bind to the licensed CD19 CARs. Affinity maturation of hits from this library led to the identification of a peptide with high affinity for the CAR. An amphiphile mimotope (amph-mimotope) vaccine based on this peptide efficiently stimulated CD19 CAR-T cells *in vitro*, triggered their expansion *in vivo*, and enhanced clearance of leukemia in both syngeneic and humanized mouse models of B-ALL/Lymphoma. To show generality, we also generated amph-mimotopes for a second human CAR recognizing the anaplastic lymphoma kinase (ALK) antigen and a murine CD19 CAR. Collectively, these results show the successful development of a clinically translatable amph-mimotope vaccine with the potential for endowing currently FDA-approved CD19 CAR-T cell therapies with enhanced anti-leukemic activity.

## Results

### Antigen^+^ tumor cells drive transient CD19 CAR-T expansion but are insufficient to promote long-term CAR-T persistence

CD19-positive relapse is often associated with limited CAR-T persistence and/or impaired functionality^3,17^. To provide insight into whether the level of available antigen stimulation impacts early CAR-T expansion/persistence, we analyzed data from several early clinical trials in children and young adults (CAYA) with B-ALL from the Children’s Hospital of Philadelphia (NCT01626495 ^18^ and NCT02906371^19^) and in adult patients with B-ALL (NCT02030847^20^) or B-cell lymphoma (NCT02030834^21^) from the University of Pennsylvania. We found that CAR-T cell expansion is generally associated with initial tumor burden, with CAR-T cells infused into patients with high tumor burden (>40% bone marrow blasts in pre-infusion bone marrow aspirate) exhibiting more pronounced expansion and longer persistence (**Fig. 1A-C, Extended Data Fig. 2A-B**). This association is statistically significant in CAYA with B-ALL (**Fig. 1A-C**) and exhibits a trend towards significance in adult B-ALL and B-cell lymphoma (**Extended Data Fig. 2A-B**), consistent with observations from earlier studies^8,9^.

**Figure 1:**
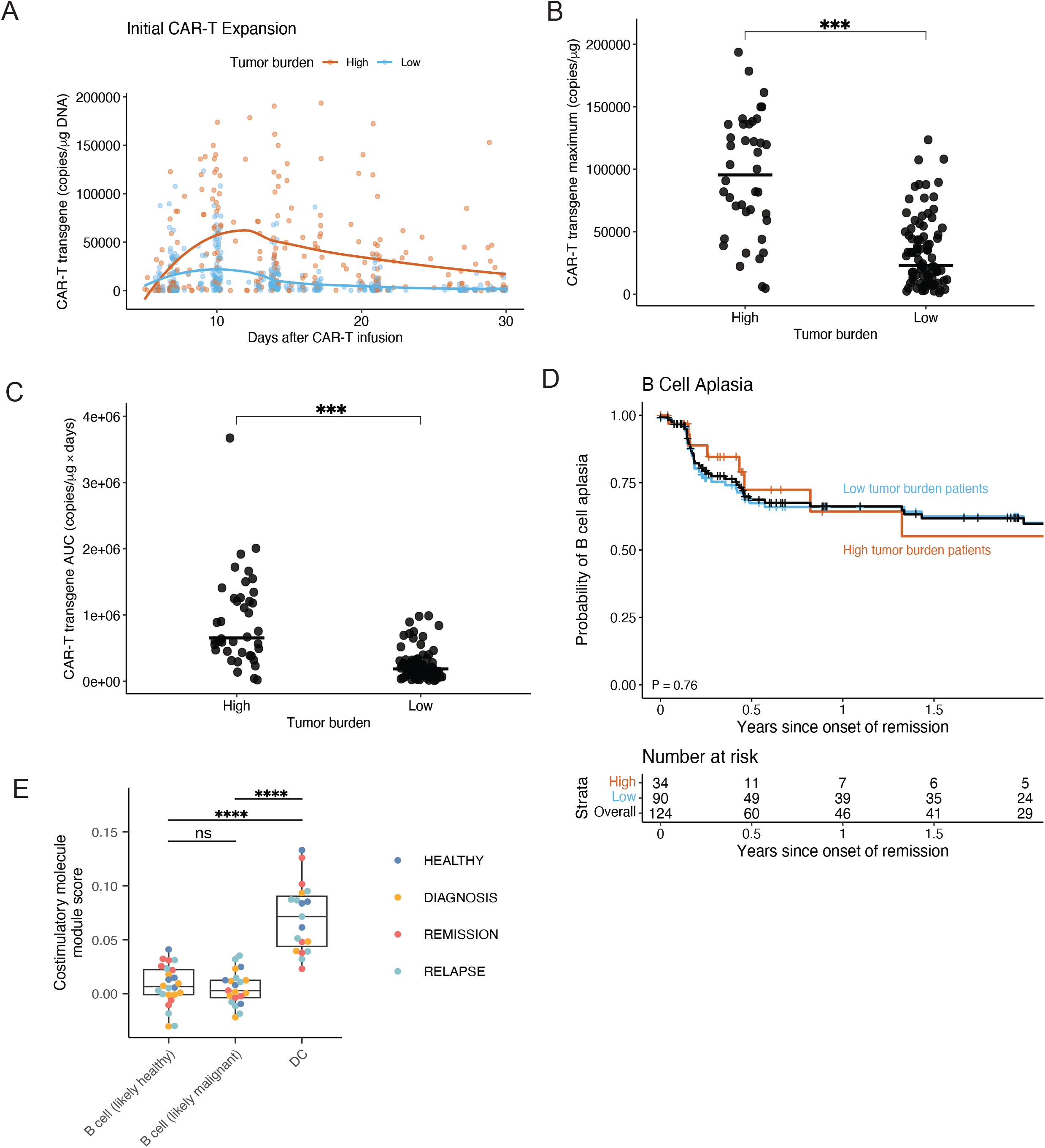
CD19 CAR-T cell expansion and B cell aplasia in pediatric B-ALL patients with initial high or low tumor burden, and rationale for a DC-targeting CD19-CAR stimulating vaccine. (**A**) Scatterplot of peripherally circulating CAR-T cells in pediatric B-ALL patients with initially high (≥40%, *n*=40) or low tumor burden (<40%, *n*=90) measured by quantitative polymerase chain reaction (qPCR) over time during the first month after infusion. Solid lines show trends generated by locally estimated scatterplot smoothing (loess). The median time to peak CAR-T expansion was 10 days after infusion in both cohorts. (**B**) Maximum concentration and (**C**) Area under the curve (AUC) of CAR-Transgene levels measured by qPCR, broken down by tumor burden. ^***^, p<0.001, by Wilcoxon rank-sum tests. (**D**) Kaplan-Meier analysis of B cell aplasia for all patients (black) or high (orange) or low (blue) tumor burden patients. B cell aplasia is defined as the time to the emergence of ≥1% CD19-positive B cell in bone marrow aspirate or ≥3% B cell by peripheral blood flow cytometry or CD19-positive relapse. Data were censored for patients who had CD19-negative relapse, reinfusion for hematogones in the marrow, alternative therapy including other CAR-T therapy or hematopoietic stem cell transplant, or non-relapse mortality. The probability of continued B-cell aplasia at two years was similar in both cohorts (p = 0.76). At 6 months, it was 72% (95% CI 55-95) in high and 67% (95% CI 57-78) in low tumor burden patients. (**E**) Box plot of costimulatory molecule module scores expressed by B cells (likely-healthy and likely-malignant) and DC. Each point represents the average of all cells of that phenotype within a patient. ^****^, p<0.0001, by a two-sided Wilcoxon rank-sum test.

Since CD19 CAR-T cells target normal B cells, continued B cell aplasia can be used as a pharmacodynamic measure for the presence of functional CD19 CAR-T cells^22^. Nearly half of the pediatric B-ALL patients receiving CD19 CAR-T cell therapy experienced B cell recovery after initial B cell aplasia (**Fig. 1D**), indicating loss of functional CAR-T cells. Interestingly, there was no significant association between the initial tumor burden and the probability of B cell recovery, suggesting that increased early stimulation from tumor cells was not sufficient to maintain CAR-T function. A recent long-term follow-up study found that a higher ratio of peak CAR-T expansion to tumor burden correlates with overall survival and is a better prediction of long-term survival than the absolute magnitude of T cell expansion or tumor burden^23^. These data suggest that if CAR-T cells receive greater antigen stimulation following infusion, they can undergo greater expansion, an important correlate of response to therapy^23,24^. However, sustained anti-tumor function over time requires that T-cells receive restimulation by professional APCs, particularly dendritic cells (DCs), which express high levels of costimulatory receptors and cytokines that are lacking from tumor cells and resting B cells^25,26^. By mining a published dataset^27^, we observed significantly higher co-stimulatory molecule expression in DCs than healthy and likely-malignant B cells in B-ALL patients (**Fig. 1E, Extended Data Fig. 3A-H**). We previously also observed that the comprehensive suite of costimulatory molecules expressed by DCs is essential for promoting memory differentiation and optimal amplification of CAR-T cells during vaccine boosting^14^. These observations suggested to us that an efficient and safe DC-targeted vaccination approach to reinvigorate and sustain CAR-T cell function could be impactful for enhancing current clinical CD19-targeting CAR-T cell therapies.

### Discovery of peptide ligands for CARs using yeast surface display

We previously demonstrated that a CAR-specific amph-vaccine could provide potent stimulation of CAR-T cells *in vivo*^14,15^. The vaccine was created by linking a CAR ligand to an albumin-binding PEG-lipid, which upon co-administration with adjuvant, would efficiently traffic to draining lymph nodes and decorate the surface of APCs, allowing presentation of the ligand to CAR-T cells in the LN (**Extended Data Fig. 1**). We demonstrated this proof of concept using CARs that recognized arbitrary small molecule ligands (e.g., fluorescein (FITC)) or linear peptide epitopes^14^. However, the antibody-based antigen-binding domain of the FDA-approved CD19 CARs (which employ an antibody clone known as FMC63) binds to a conformational epitope spanning multiple domains of the human CD19 antigen^28^. Further, the CD19 extracellular domain (ECD) is difficult to express and prone to misfolding^29^, making it challenging to use recombinant CD19 for manufacturing an amphiphile vaccine to stimulate CD19 CAR-T cells. As it is common for CAR binding domains to recognize complex conformational epitopes^30–32^, we sought to establish a general strategy to generate simple surrogate ligands for any CAR. We hypothesized that the generation of mimotopes, linear synthetic peptides that are also recognized by the CAR binding domain in addition to its native target antigen, could be an ideal option (**Fig. 2A**).

**Figure 2:**
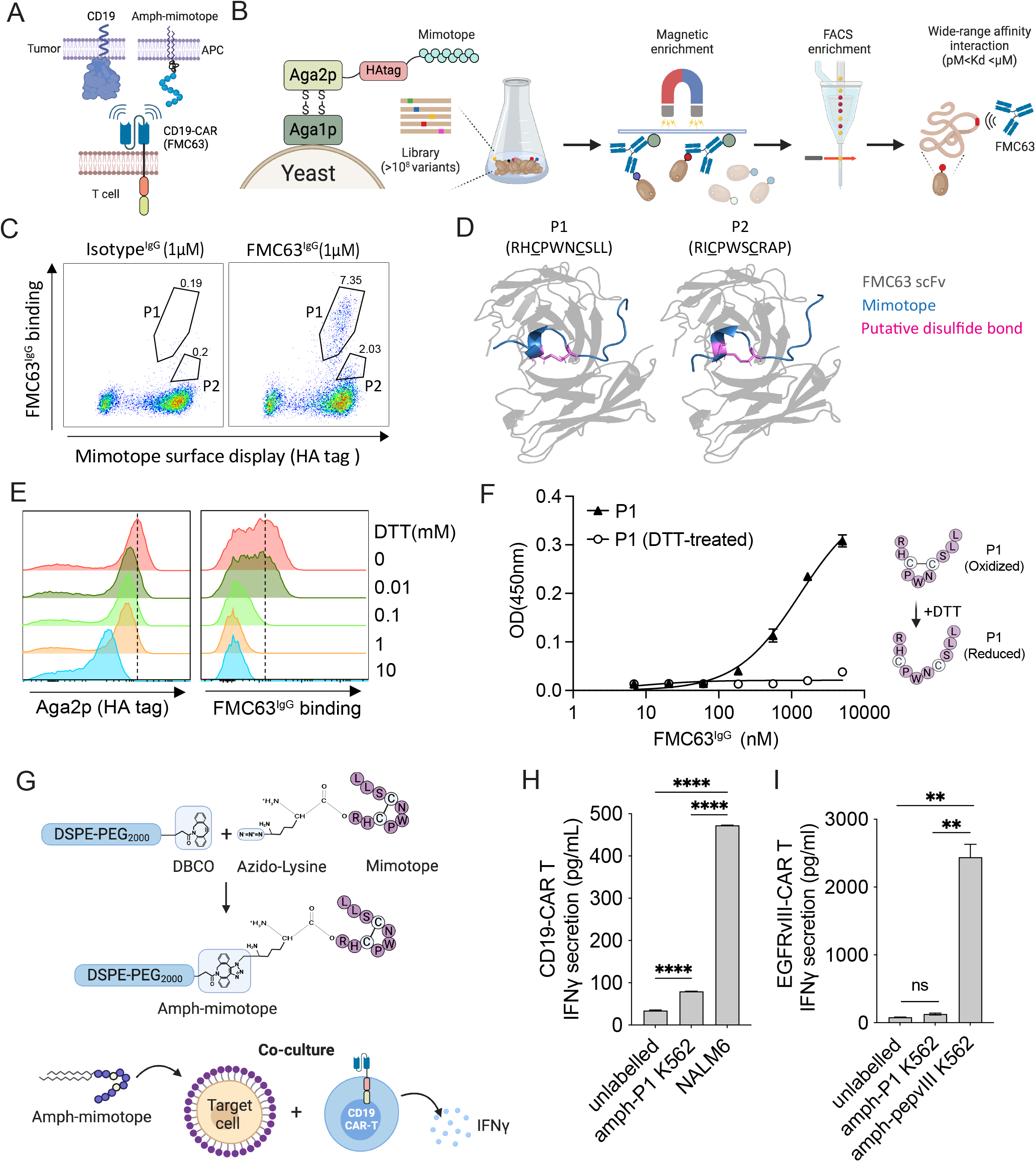
Yeast surface display identifies CD19-CAR binding mimotopes. (**A**) Illustration of CD19-CAR stimulation by natural CD19 or amph-mimotope. (**B**) Yeast surface display workflow for identifying antibody-specific mimotopes. (**C**) Flow cytometry plots showing the successful identification of yeast cell populations (P1 and P2) binding to 1 µM FMC63^IgG^ after a single round of magnetic enrichment. (**D**) Structural modeling of mimotopes P1 and P2 binding to FMC63. (**E**) Impact of the disulfide bond on mimotope binding to FMC63^IgG^ on yeast surface. Yeast cells bearing mimotope P1 were treated with increasing concentrations of DTT, and then stained with 500 nM biotin-FMC63^IgG^ and an anti-HA tag antibody for analysis of binding by flow cytometry. (**F**) ELISA analysis showing the impact of the disulfide bond on synthetic mimotope binding to FMC63^IgG^. (**G**) Scheme for amph-mimotope generation by click chemistry linkage of azide-modified P1 to DBCO-PEG-DSPE, and schematic illustration of amph-P1 coating of target cells. (**H**) CD19^-^ K562 cells labeled with 100 nM amph-P1, unlabled K562 cells, or CD19^+^ NALM6 cells were co-cultured with CD19 CAR-T at a 5:1 E:T ratio for 6 hr followed by IFN-γ ELISA. (**I**) K562 cells unlabeled, labeled with 100 nM amph-P1, or labeled with 100 nM amph-pepvIII were co-cultured with control EGFRvIII-CAR-T cells at a 5:1 E:T ratio for 6 hours followed by IFN-γ ELISA. Error bars show mean ± s.d. with triplicate samples. ^**^, p<0.01 by one-way ANOVA with Turkey’s post-test. A, B, and G are created with BioRender.com.

To identify mimotopes for FMC63, we employed a yeast surface display library^32^ presenting ∼5×10^8^ randomized linear peptides of 10 amino acids (AA, 10mer) for screening against recombinant FMC63 expressed as a full-length IgG (hereafter, FMC63^IgG^, **Fig. 2B**). Initial screening was performed using biotinylated FMC63^IgG^ attached to streptavidin beads for magnetic enrichment of weakly-bound yeast clones. After a single round of screening using this enrichment protocol, flow cytometry analysis identified two yeast populations P1 and P2, which bound with higher or lower relative affinity to FMC63^IgG^, respectively, but not an isotype control antibody (**Fig. 2C**). The two peptides shared a common enriched sequence motif RXCPWXCXXX. The presence of two cysteines in the mimotope motif suggested the possibility that these mimotopes formed an intramolecular disulfide bond, with the proline between the two cysteines likely promoting a hairpin formation in the peptide structure^33^. Computational modeling of P1 (RHCPWNCSLL) and P2 (RICPWSCRAP) interactions with FMC63 using AlphaFold predicted the mimotopes take on a curved alpha-helical structure, suggesting the formation of a cyclic structure with an intra-peptide disulfide bond (**Fig. 2D**).

To confirm the presence of a disulfide bond within the mimotope and test whether the disulfide is necessary for peptide binding to FMC63^IgG^, we treated the yeast clone P1 with increasing concentrations of the reducing agent dithiothreitol (DTT, **Fig. 2E**). Treatment with 10 mM DTT lowered Aga2p levels on the yeast cells by ∼10-fold, likely due to cleavage of the disulfide linkages anchoring the protein to the yeast cell surface. However, mild reductive conditions (i.e., 0.1mM, 1mM DTT) largely preserved Aga2p on yeast surface but eliminated FMC63^IgG^ binding (**Fig. 2E**). To further verify that this loss in antibody binding reflected loss of a disulfide in the mimotope itself, we chemically synthesized a cyclic version of mimotope P1 and monitored its binding to FMC63^IgG^ using ELISA in the presence or absence of pre-treatment of the peptide with DTT, and found that FMC63^IgG^ recognized plate-bound P1 with an apparent K_D_^IgG^ of 1.4 μM, but this binding was completely ablated for DTT-treated P1 (**Fig. 2F**). Next, we created amph-mimotope molecules by conjugating P1 functionalized with an N-terminal azide to DSPE-PEG2k-DBCO through click chemistry (**Fig. 2G**). To test whether the amph-mimotope could insert in cell membranes and enable CD19 CAR-T cell recognition, we labeled CD19-negative K562 cells with amph-P1 and then incubated them with human CD19 CAR-T cells *in vitro* (**Fig. 2G**). Amph-P1-decorated K562 cells effectively triggered IFN-γ secretion from CD19 CAR-T cells but to a much lesser extent than NALM6 cells (**Fig. 2H)**. By contrast, amph-P1 failed to stimulate control EGFRvIII CAR-T cells which responded robustly to amph-pepvIII (pepvIII, a minimal epitope from EGFRvIII, **Fig. 2I**). These results confirmed that P1 is a functional mimotope with a cyclic conformation required for its specific recognition by FMC63.

To demonstrate that the yeast mimotope library is suitable for identifying mimotopes for any desired antibody, we repeated the library screen using an anti-mouse CD19 antibody (Clone 1D3) (**Extended Data Fig. 4A-D**) and an anti-human Anaplastic Lymphoma Kinase (ALK) antibody of interest for CAR-T cell treatment of lymphoma and some lung cancers (**Extended Data Fig. 4E-F**). These additional library screens successfully identified multiple binders for each antibody, highlighting the potential of yeast surface display to effectively identify antibody-specific mimotope peptides.

### Affinity maturation of the CD19 mimotope

The affinity of a typical CAR toward its ligand is often in the low nanomolar range^34^, however, mimotope P1 only exhibited a low micromolar apparent affinity toward FMC63^IgG^, and this apparent K_D_ is likely a substantial overestimate of the monovalent affinity of the peptide for FMC63 given the bivalent IgG format used in our binding assays. We therefore aimed to further evolve the mimotope to obtain variants with enhanced affinity. To this end, a new mimotope library V2 was generated bearing a shared motif derived from the two parental mimotopes, namely RXCPWXCXXX, with a diversity of ∼1.5×10^8^ (**Fig. 3A**). We pre-enriched the V2 mimotope library for binders using FMC63^IgG^-coated magnetic beads (see Materials and Methods), and found that a significant portion of this bead-enriched library V2 could be positively stained with 20 nM biotinylated FMC63^IgG^, while the original P1 yeast clone showed no detectable binding at this concentration (**Fig. 3B**). Flow cytometry sorting of the top 1% of the positive yeast cells followed by sanger sequencing yielded dozens of mimotope clones. Sequence analysis of these mimotopes revealed a consensus sequence with high-affinity binding potential to FMC63^IgG^ (**Fig. 3C**). Several candidate mimotope clones resembling the consensus sequence were validated experimentally, and all of them exhibited stronger binding than the original clone P1 (**Fig. 3D**). We moved forward with mimotope F12 given its highest resemblance to the consensus sequence and highest binding affinity to FMC63^IgG^. This sequence was bound to FMC63 with an apparent K_D_^IgG^ of 15.6 nM (again, a value likely overestimating the true monovalent K_D_ due to the avidity effect of the bivalent FMC63 IgG). Alanine scanning across all AA positions in the F12 peptide confirmed the determining role of R at position 1°, C at positions 3° and 7°, P at position 4°, and W at position 5° (**Fig. 3E**). Residues at the remaining AA positions contributed more modestly to the overall affinity. Interestingly, inserting an extra 1-2 alanines between positions 6° and 7° completely disrupted F12 binding to FMC63^IgG^ (**Fig. 3E**), likely through the disruption of the intra-mimotope disulfide bond. Given the labile nature of the disulfide bond in the mimotope, we sought to test if the disulfide bond can be replaced with a structurally similar yet non-reducible thioacetal bond to produce a more stable mimotope (**Extended Data Fig. 5A-B**). Replacing the disulfide bond with a thioacetal bond completely eliminated its binding to FMC63^IgG^, perhaps because the thioacetal bond is 0.9 angstrom (Å) longer than the disulfide bond (**Extended Data Fig. 5C**).

**Figure 3:**
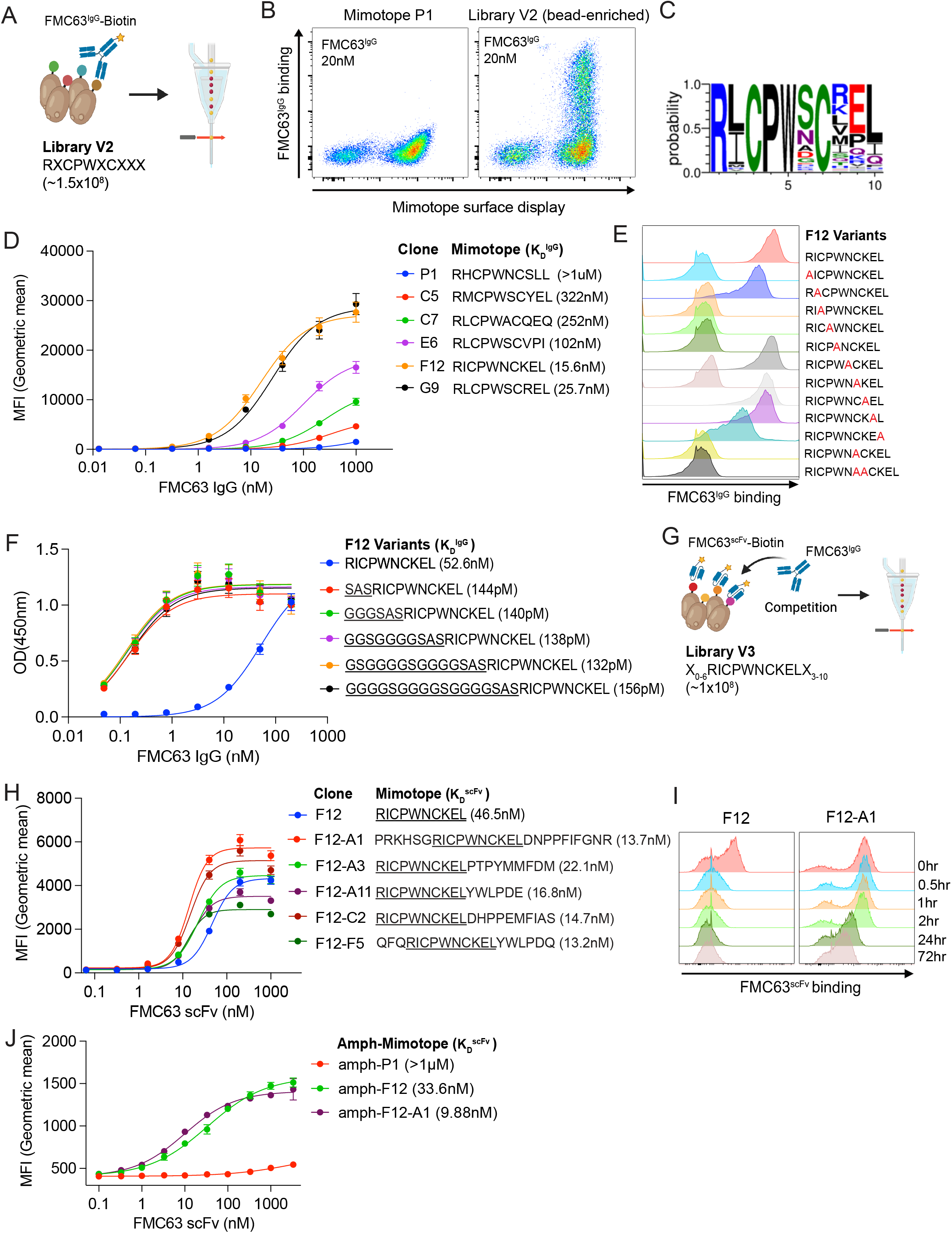
Affinity maturation of CD19 mimotopes. (**A**) Schematic workflow for identifying affinity-enhanced mimotope variants of P1 from a motif-fixed mimotope library V2. Created with BioRender.com. (**B**) Flow cytometry plots of 20 nM FMC63^IgG^ binding to yeast cells expressing the V2 mimotope library. The yeast clone P1 was included as a control. The V2 mimotope library was pre-enriched using FMC63^IgG^-coated magnetic beads (see materials and methods). (**C**) Weblogo showing the consensus mimotope sequence with high-affinity binding to FMC63^IgG^. (**D**) Binding of FMC63^IgG^ to select yeast clones assessed by flow cytometry. Amino acid sequences and apparent binding affinity are shown for each yeast clone. Median fluorescence intensities (MFI) at each concentration were fit for the calculation of apparent K_D_s. (**E**) Impact of individual AA positions on FMC63^IgG^ binding. Yeast clones bearing F12 variants with 1-2 alanine (Red) insertions or individual AA mutated to alanine (Red) were assessed for FMC63^IgG^ binding by flow cytometry. (**F**) ELISA showing the impact of flanking residues on mimotope^F12^ binding to FMC63^IgG^. The apparent binding affinity was determined as in D. (**G**) Schematic workflow for identifying affinity-enhanced mimotopes from mimotope library V3 using kinetic sorting. Yeast library V3 was first stained with monovalent biotin-FMC63scFv, followed by competition with non-biotinylated bivalent FMC63^IgG^. Created with BioRender.com. (**H**) Binding of FMC63^scFv^ to select yeast clones bearing various evolved mimotopes. The parental mimotope^F12^ sequence was underlined. The apparent binding affinity was determined as in D and specified after the mimotope sequence. (**I**) Representative histograms showing retention of prebound FMC63^scFv^ on yeast clone F12-A1 in the presence of FMC63^IgG^. Yeast cells were pre-stained with 50nM Biotin-FMC63^scFv^ followed by incubation with 1μM FMC63^IgG^. Yeast cells were sampled at indicated times for PE-streptavidin staining as an indicator for the retention of Biotin-FMC63^scFv^. (**J**) CD19^-^ K562 cells were labeled by incubation with 500 nM of the indicated amph-mimotopes, washed, then incubated with various concentrations of fluorescent FMC63^scFv^ followed by flow cytometry analysis of scFv binding; apparent binding affinities were determined as in **H**. Error bars show mean ± s.d. with triplicate samples.

Next, we sought to evaluate the contribution of the flanking sequence to mimotope binding with FMC63^IgG^. A 17 AA linker (GGGGS)_3_AS was initially introduced between the 10-mer mimotope and Aga2p during yeast library construction to avoid steric hindrance. We chemically synthesized F12 mimotope variants with increasing length of the upstream linker and performed ELISA to verify FMC63^IgG^ binding. Strikingly, the addition of even a minimal SAS tri-amino acid linker at the N-terminus of the peptide increased binding of F12 to FMC63^IgG^ by a further ∼100-fold (**Fig. 3F**). Given these data, we created a third yeast library V3 with ∼1×10^8^ diversity with 0-6 and 3-10 flanking residues extended from the N- and C-termini of F12, respectively (**Fig. 3G**), aiming to identify mimotope variants with even higher affinity. We performed kinetic sorting by competing off pre-bound biotinylated monovalent FMC63^scFv^ using non-modified bivalent FMC63^IgG^ and were able to identify yeast populations with prolonged retention of biotinylated FMC63^scFv^, indicative of slower off-rates. Characterization of individual clones identified mimotope variants with further increased affinity compared to the parental F12 clone (**Fig. 3H**). Notably, clone F12-A1 exhibited a further ∼10-fold increased affinity toward FMC63^scFv^ compared to F12. In the same competition assay as depicted in **Fig. 3G**, FMC63^scFv^ remained bound on F12-A1 yeast surface over 24 hours in the presence of excess soluble FMC63^IgG^ but was competed off the parental F12 clone within half an hour (**Fig. 3I**). Finally, we generated amphiphile conjugates of P1, F12 and F12-A1. When presented on the cell surface, these amph-mimotope variants exhibited apparent K_D_^scFv^ values of >1μM, 33.6 nM, and 9.9 nM, respectively (**Fig. 3J**). Therefore, these sequential affinity maturation screens effectively enabled the identification of mimotopes with a much-improved affinity for the CAR antibody.

### Optimized mimotopes bind to FMC63 without compromising its recognition of CD19

To gain a comprehensive view of how the CD19 mimotope interacts with FMC63, we leveraged a recently solved crystal structure of FMC63^scFv^ in complex with CD19 (PDB:7URV)^30^ and modeled mimotope binding to FMC63^scFv^. Both F12 and F12-A1 were modeled using Alphafold^35^ and Rosetta^36^, and the FMC63^scFv^-CD19, FMC63^scFv^-F12, and FMC63^scFv^-F12A1 binding interfaces were analyzed using PDBePISA^37^. PDBePISA analysis recovered many of the previously defined interactions between FMC63^scFv^ and CD19 through analysis of polarities and bond lengths **(Fig. 4A)**, confirming the validity of PDBePISA in performing interaction interface analysis. Structural modeling of F12 (**Fig. 4B**) or F12-A1 (**Fig. 4C**) interactions with FMC63^scFv^ showed that these mimotopes engage FMC63^scFv^ at the same domain. The key residues on FMC63^scFv^ that are essential for CD19 binding were found to be Y260, Y261, Y70, G263, W212, G262, S214, K69, Y265, and G129 ^30^. Of these key residues, both F12 and F12-A1 engage Y260 and S214 **(Fig. 4D)**. However, only F12-A1 engages G263 (**Fig. 4D**), which likely explains its stronger binding to FMC63^scFv^ than F12.

**Figure 4:**
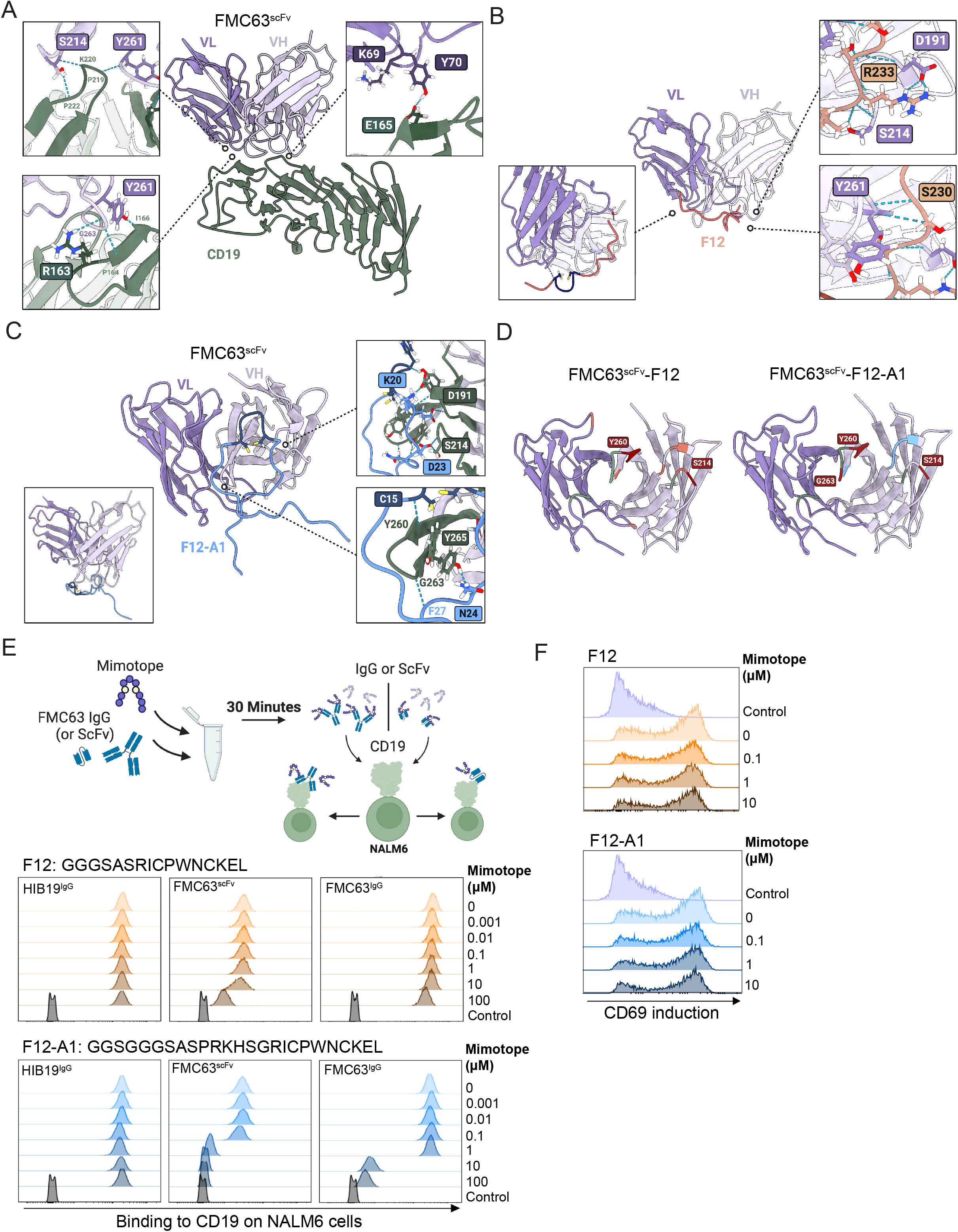
Structural modeling of mimotopes complexed with FMC63. **(A)** Ribbon diagram of FMC63^scFv^ (purple) in complex with human CD19 (green, PDB 7URV). Interactions involving key residues of FMC63 are shown. Interacting sidechain atoms are shown and boxed. Interactions involving backbone atoms are labeled without box. **(B-C)** Computational models generated using Alphafold and Rosetta for FMC63^scFv^ bound to F12 (**B**) or F12-A1 (**C**), respectively. Interactions involving previously mentioned key residues of FMC63 are labeled using the same scheme as in A. For F12-A1, the orientation of mimotope is shown to best exhibit how it engages FMC63. The panel in the bottom left depicts the complex in the same orientation as F12-FMC63. **(D)** Interface footprint of mimotopes on FMC63. On the left side, the CD19 interface is shown in green, the F12 interface shown in orange, and overlapping residues are shown in red. On the right side, the CD19 interface is shown in green, the F12A1 interface shown in blue, and overlapping are residues shown in red. **(E)** Recognition of natural CD19 recognition by FMC63^scFv^ in the presence of mimotopes. Biotinylated FMC63^scFv^ was pre-incubated with various concentrations of F12 or F12-A1 prior to staining NALM6 cells and detection via PE-conjugated streptavidin. For FMC63^IgG^ binding, NALM6 cells were stained with the same concentrations of mimotope and FMC63^IgG^ with no prior incubation, and binding was detected using PE-conjugated anti-mouse-IgG. A non-competing IgG clone HIB19 is shown as a control. **(F)** Activation of mimotope-bound CD19 CAR-T cells by target cells. CD19-CAR Jurkat T cells were pre-incubated with F12 or F12-A1 at indicated concentrations, then co-cultured with NALM6 cells at a 1:1 E:T ratio for 16 hours followed by flow cytometry analysis of CD69 expression. Shown in E-F are one representative results of three independent experiments.

To experimentally assess potential overlaps in the binding interface of FMC63 to CD19 vs. our mimotope peptides, we pre-blocked FMC63^scFv^, FMC63^IgG^ or an alternative control anti-CD19 antibody HIB19^IgG^ with increasing concentrations of mimotopes F12 or F12-A1, and then added the antibodies to CD19^+^ NALM6 cells (**Fig. 4E**). When FMC63^scFv^ was pre-blocked with F12 peptide, binding to NALM6 cells was only compromised when the F12 concentration was 100 µM, and NALM6 binding was nearly unaffected for FMC63^IgG^. By contrast, F12-A1 abolished FMC63^scFv^ or FMC63^IgG^ binding to CD19 when the mimotope concentrations were greater than 1 or 10 µM, respectively, consistent with its higher affinity for FMC63. However, when FMC63 CAR-expressing Jurkat T cells were pre-blocked with increasing concentrations of F12 or F12-A1 and stimulated with NALM6 cells, CD69 induction on Jurkat cells was not affected (**Fig. 4F**). Thus, despite some ability of the mimotopes to interfere with soluble FMC63 binding to CD19^+^ cells at high concentrations, FMC63-CAR-T cells retained their full potential of recognizing CD19^+^ leukemic cells even in the presence of mimotopes.

### Amph-mimotope vaccination triggers FMC63-mCAR-T expansion *in vivo*

To test if amph-mimotope vaccines could efficiently stimulate human CD19-targeting CAR-T cells in an immunocompetent mouse model *in vivo*, we established a hybrid CAR by fusing the FMC63 scFv to murine CAR signaling domains, (FMC63-mCAR, **Fig. 5A**). First, we validated the ability of the amph-mimotope variants to stimulate FMC63-mCAR-T cells. K562 cells were pre-labeled with amph-mimotope variants and co-cultured with FMC63-mCAR-T cells as in **Fig. 2G**. Amph-mimotope-decorated target cells triggered IFN-γ production from the CAR-T cells, with the stimulation efficiency positively correlating with FMC63-mimotope affinity and amph-F12-A1 providing the strongest CAR-T activation, comparable to stimulation by NALM6 cells (**Fig. 5B**). To determine if these amph-mimotopes could be effectively presented by DCs *in vivo*, we vaccinated C57BL/6 mice with 10 nmol amph-F12-A1, a dose we previously found to be effective for amph-peptide stimulation of CAR-T cells. We formulated the mimotope with or without cyclic-di-GMP, a STING agonist added as an adjuvant^38^. Following vaccination, we isolated CD11b^+^ myeloid cells and CD11c^+^ DCs from the draining inguinal lymph nodes and analyzed the presence of surface-displayed amph-peptides using flow cytometry by staining the cells with biotinylated FMC63^IgG^ (**Extended Data Fig. 6A**). Interestingly, high levels of amph-mimotope were detected on APCs for at least 24 hr, but only when amph-mimotope was co-administered with adjuvant (**Fig. 5C**).

**Figure 5:**
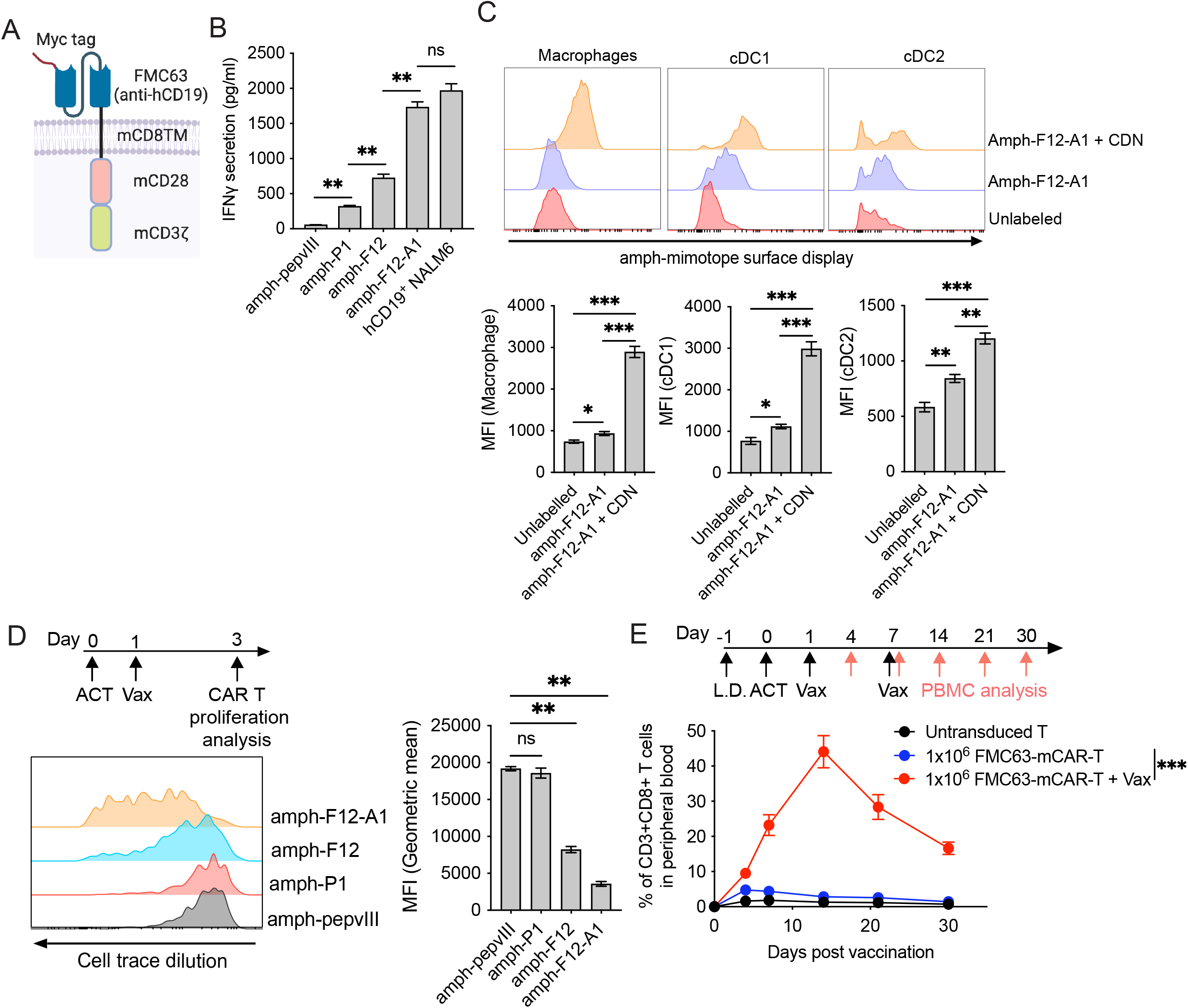
Amph-mimotopes require a threshold affinity to stimulate CAR-T proliferation in vivo. (**A**) Schematic of hybrid FMC63-mCAR design. (**B**) Comparison with various amph-mimotopes in stimulating FMC63-mCAR-T cells. K562 cells were labeled with 100 nM of amph-mimotope variants for 30 min, washed, and co-cultured with FMC63-mCAR-T cells at a 1:1 E:T ratio for 6 hours followed by ELISA measurement of IFN-γ secretion. (**C**) Amph-mimotope labeling of APCs *in vivo*. C57BL/6 mice (*n*=3 animals/group) were vaccinated with 10 nmol amph-mimotope ± CDG adjuvant, and 24 or 48 hr later, mimotope uptake by macrophages and cDCs was detected by staining with FMC63 followed by flow cytometry analysis. Shown are representative histograms of FMC63 staining. (**D**) Amph-mimotope vaccine stimulation of CAR-T cells *in vivo*. WT C57BL/6 mice (*n*=3 animals/group) received adoptively transferred with 2×10^6^ CTV-labeled FMC63-mCAR-T cells, then vaccinated 1 day later with 10 nmol amph-peptides + 25μg CDG adjuvant (Vax). Shown above is the timeline and below are representative histograms of FMC63-mCAR-T cell proliferation in LNs 48 hours after vaccination. (**E**) Longitudinal monitoring of CAR-T expansion in response to amph-mimotope vaccination. C57BL/6 mice (*n*=5 animals/group) were lymphodepleted (L.D.), adoptively transferred with 10^6^ FMC63 CAR-T cells, and then vaccinated at indicated time points, and circulating FMC63-mCAR-T cells were quantified in the blood by flow cytometry over time. Error bars show mean ± s.d. with triplicate samples for B, mean ± 95% CI for C-E. ^***^, p<0.0001;^**^, p<0.01; n.s., not significant, by one-way ANOVA with Turkey’s post-test for B-D, two-way ANOVA with Tukey’s post-test for E.

Next, we labeled FMC63-mCAR-T cells with cell trace violet (CTV) and transferred them into C57BL/6 mice, followed by vaccination with amph-P1, amph-F12, amph-F12-A1 vaccine, or an irrelevant amph-pepvIII as a control and flow cytometry analysis to detect CAR-T cell proliferation (CTV dilution) 2 days later. This functional test of amph-mimotope stimulation revealed a clear hierarchy of mimotope activity *in vivo*, with amph-P1 eliciting no CAR-T cell proliferation, amph-F12 eliciting limited proliferation, and only amph-F12-A1 triggering proliferation of the entire CAR-T population (**Fig. 5D**). These data suggest there is an affinity threshold for mimotope vaccines to effectively stimulate CAR-T cells *in vivo*.

Clinical evidence indicates that lack of antigen exposure soon after adoptive transfer may compromise CAR-T engraftment^39^. To test if amph-mimotope vaccine boosting post adoptive cell transfer could promote FMC63-mCAR-T cell expansion and engraftment in the absence of leukemia, naïve CD45.2^+^ C57BL/6 mice received sublethal lymphodepletion followed by intravenous infusion of either CD45.1^+^ FMC63-mCAR-T or control untransduced T cells (**Fig. 5E**). Without additional treatment, peripheral CD45.1^+^ CAR-T cells only occupied <5% of total CD8^+^ T cells by day 4 and decreased to <3% by day 14 (**Fig. 5E**). By contrast, amph-F12-A1 vaccine boosting rapidly amplified CD45.1 CAR-T cells, reaching ∼40% of total peripheral CD8^+^ T cells (∼500 CAR-T cells per microliter of blood, **Fig. 5E, Extended Data Fig. 6B**) by day 14 after two weekly vaccinations, and then slowly contracting over the next few weeks.

### Amph-mimotope vaccine boosting of FMC63-mCAR-T in an immunocompetent mouse model of B-ALL/Lymphoma

We next sought to evaluate effects of optimized amph-F12-A1 vaccine boosting of FMC63 CAR-T in an immunocompetent mouse model of CD19^+^ hematological malignancy. We transduced Eμ-Myc B-ALL/Lymphoma cells^40^ with human CD19 and firefly luciferase. When these hCD19^+^Luc^+^ Eμ-Myc cells were inoculated into C57BL/6 mice, animals developed aggressive B-ALL/Lymphoma with 100% penetrance and a median survival of 18 days (**Extended Data Fig. 7A-C**). FMC63^IgG^ showed minimal binding WT B cells expressing mouse CD19, and FMC63-mCAR-T effectively killed hCD19^+^ but not WT Eμ-Myc cells *in vitro* (**Extended Data Fig. 7D-E**). On day 4 post leukemia engraftment, mice were infused with FMC63-mCAR-T alone of CAR-T combined with three weekly amph-mimotope vaccine boosts (**Fig. 6A**). Treatment with CAR-T alone resulted in moderate leukemia control (**Fig. 6A-B**); however, CAR-T in combination with amph-mimotope boosting markedly suppressed disease progression (**Fig. 6A-B**). Enumeration of CAR-T cells in the blood confirmed that vaccine boosting elicited a rapid CAR-T expansion (**Fig. 6C**). Phenotypically, peripheral CAR-T cells from mice receiving vaccination were dominated by an effector population on day 11, but by day 18, nearly 50% of CAR-T cells in vaccinated mice exhibited a central memory phenotype compared to <20% in non-boosted animals (**Fig. 6D-E**). Vaccine-boosted CAR-T cells also possessed significantly higher cytokine polyfunctionality (**Fig. 6F-G**). As a result, mice receiving both CAR-T and amph-mimotope vaccine had greatly extended survival compared to those receiving CAR-T cells alone (**Fig. 6H**). Lymph nodes of amph-mimotope-boosted animals retained organized T and B cell areas suggesting a lack of overt toxicity of the vaccine boost to draining lymph nodes (**Extended Data Fig. 8A-B**), and no mimotope-specific antibodies were induced (**Extended Data Fig. 8C**). Thus, amph-mimotope boosting of FMC63 CAR-T cells induced multiple favorable effects on CAR-T cell phenotype and increased the anti-tumor efficacy of CAR-T therapy.

**Figure 6:**
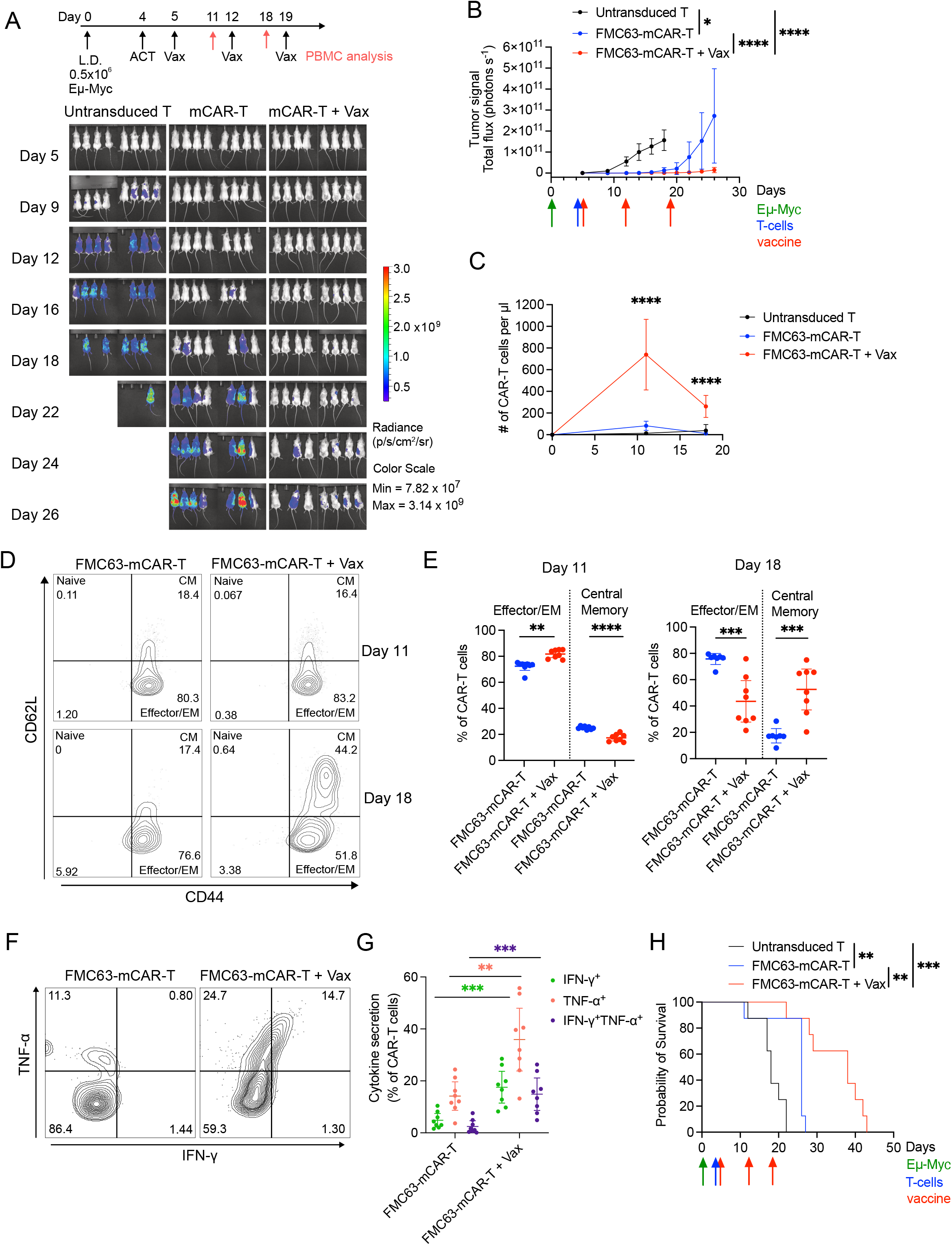
Amph-mimotope vaccine enhances FMC63-mCAR-T therapy in an immunocompetent hCD19^+^ Eμ-Myc B-ALL/Lymphoma mouse model. (**A-B**) Real-time whole animal imaging tracking disease progression. C57BL/6 albino mice were lymphodepleted (L.D.) and injected with 0.5×10^6^ hCD19^+^ Luc^+^ Eμ-Myc cells. On day 4, mice were adoptively transferred with 2×10^6^ FMC63-mCAR-T cells, then vaccinated 1 day later with 10 μg amph-F12-A1 + 25 μg CDG(*n*=8 animals/group). Shown are whole animal imaging at indicated times (**A**), and quantification of total photon counts over time in each group (**B**). (**C**) Enumeration of circulating CAR-T cells or control T cells by flow cytometry over time. (**D-E**) Immunophenotype of circulating CAR-T cells on days 11 and 18. Shown are the representative flow cytometry plots (**D**), and percentages of effector/effector memory (EM, CD44^+^CD62L^-^) and central memory (CM, CD44^+^CD62L^+^) CAR-T cells (**E**). (**F-G**) Cytokine polyfunctionality of circulating CAR-T cells on day 11. Shown are representative contour plots (**F**), and percentages of cytokine-secreting CAR T cells (**F**). (**H)** The overall survival of Eμ-Myc mice under various treatment regimens. Error bars show mean± 95%CI. ^*^, p=0.0100; ^**^, p<0.01; ^***^, p<0.001, ^****^, p<0.0001, by two-way ANOVA with Tukey’s post-test for B, by one-way ANOVA with Tukey’s post-test for C, by unpaired t-test for E, G, by Log-rank (Mantel-Cox) test for H.

We previously reported amphiphile-fluorescein (amph-FITC) boosting of CAR-T cells bearing bivalent FITC/tumor antigen-specific CARs^14^. To compare amph-mimotope boosting of the FMC63-mCAR vs. this alternative approach in the hCD19^+^ Eµ-Myc model, we synthesized a tandem FITC/hCD19-targeted murine CAR construct (i.e., αFITC-FMC63-mCAR), employing the same high-affinity anti-FITC scFv we previously tested. *In vitro*, these dual-CAR T cells effectively recognized and killed hCD19^+^ Eµ-Myc cells (**Extended Data Fig. 7E**). Treatment of Eµ-Myc tumor-bearing animals with tandem αFITC-FMC63-mCAR-T cells alone elicited similar tumor control as the monovalent FMC63-mCAR-T (**Extended Data Fig. 9A-C**, compare with **Fig. 6A-B**). However, while animals receiving treatment with the tandem CAR-T alone did not show signs of toxicity until the tumors progressed, all animals receiving the amph-FITC vaccine boost rapidly developed signs of severe toxicity immediately following the first vaccine boost, including hypothermia and hunched posture, and despite similar early tumor control to the non-boosted tandem CAR group, all animals receiving amph-FITC boosting died by day 20 (**Extended Data Fig. 9A-D**). By contrast, mice receiving FMC63-mCAR-T + the amph-mimotope vaccine showed no visible signs of toxicity during the treatment course (**Extended Data Fig. 9D**).

As the major difference between the amph-FITC and amph-mimotope vaccines is the affinity of the ligands for the CAR (the 4m5.3 scFv used in the FITC CAR binds to fluorescein with K_D_ = 270fM^41^), we hypothesized that these disparate outcomes reflect an over-stimulation of the tandem αFITC-FMC63-mCAR T cells, leading to an exacerbation of cytokine release syndrome (CRS) when these cells encounter circulating tumor cells in the B-ALL/Lymphoma model. Comparison of CAR-T cell levels in the blood showed that CAR-T expansion induced by the two vaccines was comparable, suggesting that increased toxicity following FITC boosting could not be attributed simply to the number of CAR-T cells present (**Extended Data Fig. 9E**). However, analysis of serum cytokines at day 6 after the first immunization (or earlier from moribund mice) showed severe CRS in mice receiving amph-FITC vaccine + tandem CAR-T as exemplified by massive elevations of serum IL-6, IL-10, and TNF-α (**Extended Data Fig. 9F**). However, these signature CRS cytokines remained at low levels in mice receiving CAR-T only, or FMC63-mCAR-T plus amph-mimotope vaccination (**Extended Data Fig. 9F**). To assess if this discrepancy was triggered by the differential responses of CAR-T cells towards their respective vaccines, we co-cultured CAR-T cells with the amph-vax-labeled K562 cells *in vitro* and found that amph-FITC triggered much higher levels of IFN-γ and TNF-α secretion by tandem CAR-T cells than the amph-mimotope elicited cytokine secretion from FMC63-mCAR-T cells (**Extended Data Fig. 9G-H**). These results suggest that in the setting of cancers where CAR-T cells encounter a substantial frequency of tumor cells in the blood, high-affinity ligand boosting can exacerbate CRS. This provides an important additional rationale for the mimotope discovery pipeline, where moderate affinity ligands can be readily isolated for any given CAR.

### Amph-mimotope-labeled DCs enhance human CD19 CAR-T expansion and leukemia clearance in a humanized mouse model

Human CAR-T cells are routinely evaluated in immunodeficient NSG mice, but this animal model is problematic for testing lymphatics-mediated amph-vaccine boosting, as these mice have generally defective lymph nodes and lymphatic development due to their lack of native lymphocytes^42^. To overcome this technical hurdle, we devised an alternate strategy to test amph-vaccine boosting of human CAR-T cells. We previously showed that amph-ligand vaccine molecules label lymph node-resident DCs, which are the key APC population mediating the amph-vaccine stimulation^14^. To mimic this process in NSG mice in a human-relevant setting, we differentiated human peripheral blood monocytes into DCs *in vitro* and matured them with lipopolysaccharide (LPS) and IFN-γ (**Fig. 7A**). These matured human monocyte-derived DCs (MoDCs) exhibited significantly increased expression of multiple co-stimulatory molecules, including CD80/86, 41BBL, OX40L, and ICOSL (**Fig. 7A**). Incubation of activated MoDCs with amph-F12-A1 led to robust cell surface labeling with the mimotope (DC-mVax, **Fig. 7B**). DC-mVax efficiently stimulated human CD19 CAR-T cell activation *in vitro* (**Fig. 7C**) but were refractory to CAR-T cell killing (**Extended Data Fig. 10A**), consistent with our prior observations that DCs were not eliminated during vaccine boosting *in vivo*^14^, and work suggesting DCs have mechanisms to resist CTL-mediated killing^43,44^. To determine if DC-mVax could activate and expand human CD19 CAR-T cells *in vivo* in the absence of additional antigen, we infused human CD19 CAR-T cells into non-leukemic NSG mice with or without subsequent *i.v*. infusion of DC-mVax (**Fig. 7D-E**). Mice receiving a single dose of DC-mVax 24 hours post CAR-T infusion exhibited significant expansion of CAR-T cells in the peripheral blood compared to those receiving unmodified DCs, and circulating human CD19 CAR-T cells remained detectable at day 30 (**Fig. 7E**). Notably, the extent of CAR-T cell expansion markedly decreased when DC-mVax was infused 7 days post CAR-T cell transfer, indicating that early antigen exposure plays an important role in promoting CAR-T expansion and engraftment (**Fig. 7E**). Repeated dosing of DC-mVax on day 1 and day 7 days led to a further increase in CAR-T cell expansion (**Fig. 7E**).

**Figure 7:**
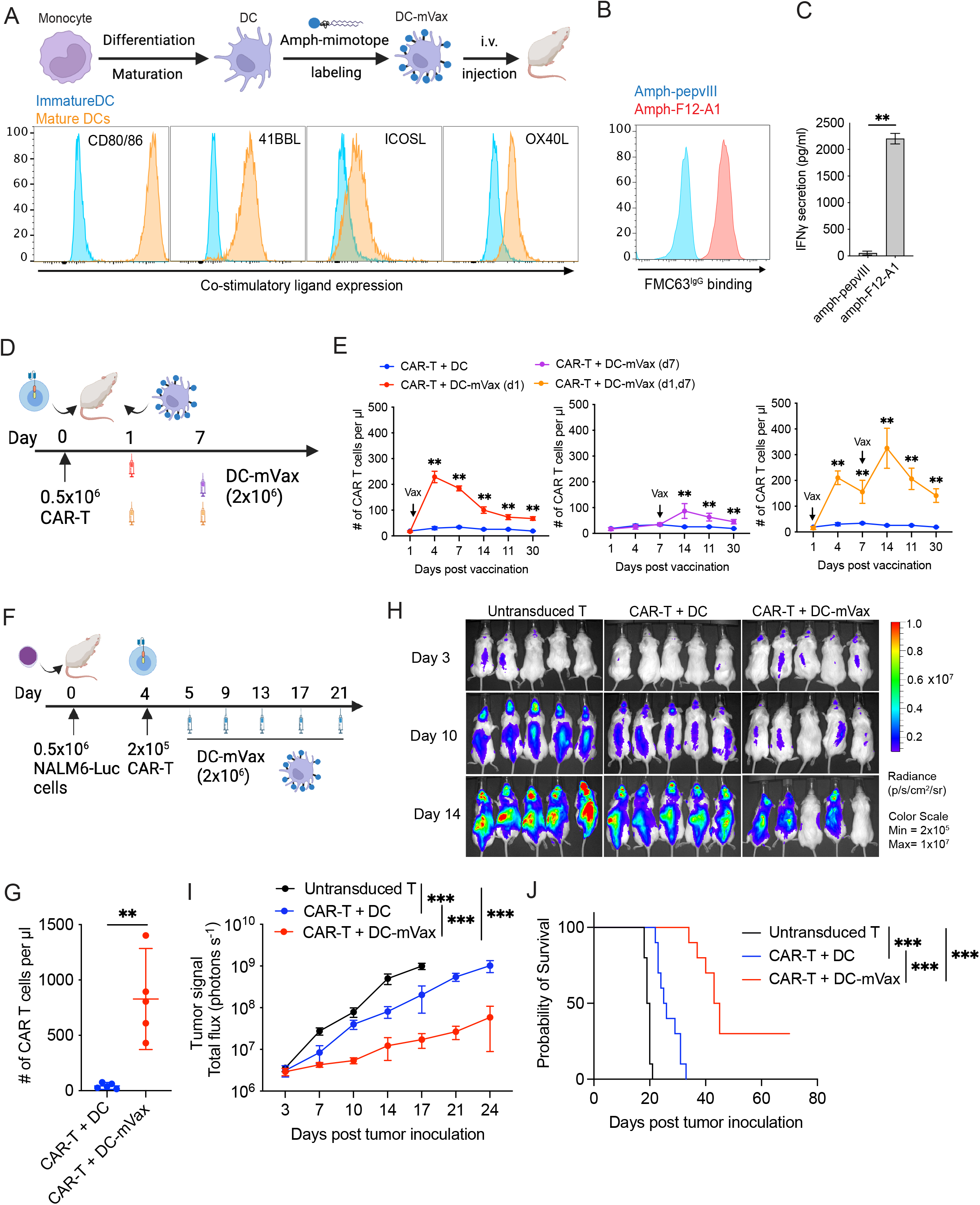
Amph-mimotope vaccine-labeled DCs augment human CD19 CAR-T therapy against leukemia. (**A**) Schematic of approach to generate “DC-mVax” and representative histograms depicting LPS/IFN-γ-induced upregulation of co-stimulatory markers on mature DCs. (**B**) Representative histogram showing FMC63^IgG^ binding to DC-mVax. Activated MoDCs were incubated with 500 nM amph-F12-A1 or control amph-pepvIII for 30min, washed, and then stained with FMC63^IgG^. (**C**) CAR-T activation by DC-mVax. Activated MoDCs were labeled with amph-F12-A1 or amph-pepvIII as in (**B**) and then co-cultured human CD19 CAR-T cells at a 1:1 E:T ratio. Shown is IFN-γ measured in the supernatant after 6 hr by ELISA. (**D**) Experimental setup and timeline for E. (**E**) Quantification of circulating human CD19 CAR-T cells in the absence or presence of a single dose of DC-mVax at day 1 or day 7 or two weekly doses of DC-mVax administered following timeline in D (n=5 animals/group). (**F**) Experimental setup and timeline for G-J. (**G-J**) Real-time whole animal imaging of B-ALL progression in NSG mice. Human CD19-CAR-T cells were administered with or without DC-mVax and tumor progression was monitored using *in vivo* bioluminescent imaging (*n*=5 animals/group. Shown are the enumeration of circulating human CD19 CAR-T cells on day 21 (**G**), whole animal imaging at indicated times (**H**) and quantification of total photon counts over time (**I**). Shown is one of two independent experiments. (**J**) Overall survival of NSG mice under various treatment regimens. *n*=10 animals assembled from two independent experiments. Shown are combined data from two independent experiments. Error bars show mean ± s.d. with triplicate samples for C, and mean ± 95% CI for E,G, I-J. ^***^, p<0.0001; ^**^, p<0.01 by Student’s t-test for C and G, by two-way ANOVA with Tukey’s post-test for E and I, and by Log-rank (Mantel-Cox) test for J.

To determine if DC-mVax boosting could augment the anti-tumor activity of human CD19 CAR-T cells, we established leukemia in NSG mice by infusing luciferase-expressing NALM6 tumor cells as previously described^45^. Suboptimal doses of CD19 CAR-T cells were administered on day 4 followed 24 hours later by repeated *i.v*. infusions of either unmodified DCs or DC-mVax (**Fig. 7F**). Despite the presence of high levels of target cells, CAR-T cells alone showed poor persistence/expansion compared to CAR-T cells boosted with DC-mVax (**Fig. 7G**). Notably, DC-mVax boosting significantly enhanced CAR-T cytokine polyfunctionality (**Extended Data Fig. 10B-C**) and differentiation toward a central memory phenotype (**Extended Data Fig. 10D-E**). While CAR-T cells alone only slightly delayed tumor expansion and moderately prolonged survival, the addition of the DC-mVax boosting enabled this dose of CAR-T cells to substantially slow tumor growth (**Fig. 7H-I**) and 30% of the animals completely rejected their tumors (**Fig. 7J**). Collectively, these results demonstrated that a suitably affinity-matured amph-mimotope could act as an effective vaccine to promote engraftment, expansion, and reinvigoration of human T cells expressing the CAR currently used in all approved CD19-targeting CAR-T products and enhance their anti-leukemic activity.

## Discussion

Successful engraftment and persistence is critical for ensuring robust and long-term anti-leukemic activity by CAR-T cells^1,46^. Analysis of data from recent CAR-T cell clinical trials suggested that CD19-directed CAR-T cells initially expand more strongly if there is a high tumor burden, consistent with the idea of CAR-T expansion that is commensurate with the dose of antigen encountered. However, the long-term persistence of CAR-T cells does not correlate with the initial tumor burden, suggesting that this initial stimulus vanishes over time as leukemic cells are cleared from the body and/or that stimulation received from tumor cells or naïve B cells lacking appropriate costimulatory signals is insufficient to maintain the CAR-T cell population. We and others have recently demonstrated the ability of using a synthetic vaccine to boost CAR-T cells *in vivo* and achieved enhanced therapeutic outcomes^14,15,47^. Our strategy of linking a ligand for the CAR to a lymph node-targeting PEG-lipid amphiphile is simple but requires a suitable peptide or protein ligand. The poor expression and misfolding of recombinant human CD19 ectodomain^29^ make the use of the native target antigen challenging for this approach. Here we demonstrate that surrogate CAR ligands based on short peptides can be readily identified and affinity-matured using directed evolution, enabling the generation of mimotope vaccines that efficiently boost CD19 CAR-T cells analogous to natural CD19 antigen exposure. Vaccine boosting significantly impacted the outcome of CD19 CAR-T cell therapy by increasing CAR-T cell expansion and engraftment as well as augmenting CAR-T cell functionality. Importantly, the mimotope vaccine discovered here is applicable to all four FDA-approved FMC63-based CD19 CAR-T cell therapies.

In addition to the human CD19 CAR mimotope, mimotopes for anti-mouse CD19 and anti-human ALK CARs were also successfully identified using the same yeast mimotope library, indicating the broad applicability of this library for identifying mimotopes for any desired antibody. Experimental verification and structural modeling demonstrated that the CD19 mimotope discovered through the yeast library forms a cyclic conformation that is essential for its recognition by FMC63. Given that FMC63 recognizes a conformational epitope, it is possible that a mimotope also needs to form a desired secondary structure to achieve strong binding. Notably, the CD19 mimotope only partially occupies the natural epitope binding domain on FMC63. This partial occupancy appears to enable FMC63-based CD19 CAR-T cells to maintain their capacity to recognize CD19^+^ leukemia cells even in the presence of the mimotope, which could be important for allowing the CAR-T cells to continue sensing tumor cells in lymph nodes during vaccine boosting. The transient surface presentation of amph-ligands also prevents CAR-T cells from receiving excessive and prolonged antigen exposure, thus reducing the likelihood of CAR-T cell exhaustion and severe cytokine release syndrome^14^.

We previously reported an alternative approach for vaccine boosting CAR-T, where the CAR itself is engineered as a tandem CAR with one binding domain recognizing a tumor antigen and the other domain binding to the synthetic molecule FITC, enabling boosting of these tandem CARs with the amph-FITC vaccine^14,15^. While this approach avoids the need to generate a new amph-vaccine for each new CAR product, many CARs (such as the FMC63-based CAR products) have already reached extensive clinical testing, providing a strong motivation to leave the CAR design itself unchanged. In addition, the tendency of scFvs to undergo aggregation that can induce detrimental tonic signaling in CAR T cells^48–50^ may make the bivalent CAR approach untenable for some CARs. In these settings, the amph-mimotope strategy is an ideal solution.

While B cells retain the antigen for CAR-T, evidence suggests that B cells themselves may not be as effective as conventional DCs for restimulating CAR-T cells at least partially due to greater expression of co-stimulatory molecules in DCs^26^. Our previous findings showed that co-stimulatory molecules expressed by activated DCs are crucial for optimally expanding CAR-T cells with amph-ligands^14^. This DC-specific effect is further exemplified in our DC-mVax boosting of human CAR-T in NSG mice and amph-mimotope vaccine boosting of mouse CAR-T in the Eμ-Myc B-ALL/Lymphoma mouse model, where vaccination not only drove CAR-T expansion but also enhanced effector cytokine expression and promoted memory differentiation. Conventional CD19 CAR-T cells expand and show efficacy following the first infusion, however, reinfusion of CAR-T cells into patients experiencing CD19^+^ tumor relapse is often ineffective, with significantly lower CAR-T expansion compared to the first infusion^39,51–53^. Therefore, one potential use of the mimotope vaccine is to support CAR-T cells during reinfusion. Notably, even during the first infusion, CD19 CAR-T cells rapidly decline after a transient expansion^9,54^. Despite the general correlation of initial CAR-T expansion with baseline tumor burden, a long-term follow-up study in B-ALL found that a higher ratio of peak CAR-T-cell expansion to tumor burden significantly correlated with event-free survival and overall survival^23^. Although this effective ratio more likely occurs in patients with lower tumor burden, administering the mimotope vaccine upon CAR-T contraction during the first infusion could potentially reinvigorate CAR-T cells and sustain CAR-T expansion to elevate this effective ratio in patients with both low and high tumor burden to improve event-free survival.

Given the toxicity associated with rapid CAR-T activation and expansion in patients with hematological malignancies, such as CRS and immune effector cell-associated neurotoxicity syndrome (ICANS), amph-vaccine affinity, dose, and dosing intervals may need to be tailored based on the CAR-T cell number and residual tumor burden in patients in future clinical trials. In our previous work, the amph-FITC vax/tandem CAR-T combination enhanced CAR-T therapy against solid tumors with transient mild toxicity^14^. However, in a side-by-side comparison, we found that the amph-FITC vax/tandem CAR-T combination triggered a lethal CRS in the immunocompetent Eμ-Myc B-ALL/Lymphoma mouse model, while amph-mimotope vax/CAR-T treatment effectively controlled leukemia progression with limited toxicity. Our data suggests this drastic discrepancy in toxicity profiles is linked to the much higher affinity of the tandem CAR towards the amph-FITC vaccine vs. the amph-mimotope binding to FMC63. These data indicate that amph-vaccine ligand affinity plays a critical role in the outcome of CAR-T boosting in cancers involving substantial levels of circulating tumor cells that will trigger cytokine secretion in the blood. The mimotope discovery pipeline described here is well suited to identify effective and safe ligands, as it allows mimotopes with a wide range of affinities to be discovered.

One set of challenges for characterizing amph-vaccines designed for human T cells are the deficiencies of current humanized mouse models. The defective lymphatics, absent or disorganized lymph nodes, and poor crosstalk of mouse APCs with human T cells is a barrier to evaluating vaccine boosting of human CAR-T cells in NSG mice^42^. As an alternative, we evaluated vaccine boosting of mouse T cells bearing a hybrid FMC63-mCAR in immunocompetent C57BL/6 mice via standard vaccine administration routes and boosting of human CD19 CAR-T cells using mimotope-decorated human MoDCs in NSG mice via *i.v*. delivery. These two complementary approaches demonstrated the feasibility of the mimotope vaccine in the setting of natural trafficking and distribution from lymphatics to lymph nodes, as well as its capacity to stimulate human CAR-T cells upon insertion into the surface of human DCs.

mRNA vaccines encoding the target tumor antigen can also be employed to stimulate CAR-T cells^47^. Recently, a CLDN6-expressing mRNA vaccine was tested in a phase 1 clinical trial in combination with CLDN6-directed CAR-T cells and showed promise in improving CAR-T cell expansion in patients with solid tumors^55^. Importantly, CAR-T plus mRNA vaccination was well tolerated, suggesting the overall safety profile associated with this combination therapy. However, mRNA-LNPs differ from our amphiphile vaccines in their *in vivo* trafficking, biodistribution, and the duration of antigen presentation on the cell surface. Therefore their differential effects on boosting human CD19 CAR-T cells remain to be determined.

In summary, we have designed and validated a general strategy to generate mimotope ligands for any CAR. Using this methodology, we demonstrate an amph-mimotope vaccine that could be used clinically in tandem with all four FDA-approved CD19 CAR-T cell products. The ability of amph-mimotope vaccines to efficiently stimulate human CD19 CAR-T cells *in vivo* supports its potential use as a robust and clinically translatable approach for enhancing the engraftment, long-term persistence, and anti-leukemic activity of current CD19 CAR-T cell therapies.

## ACKNOWLEDGMENTS

We thank the Koch Institute Swanson Biotechnology Center for technical support, specifically the flow cytometry core facility. We thank Dr. Michael Hemann from the Koch Institute of MIT for sharing the Eμ-Myc cells. We thank Dr. Steve Albelda for his suggestions on the manuscript.

## Data and materials availability

All data are available on request.

## AUTHOR CONTRIBUTIONS

L.M., D.J.I., and K.D.W designed the studies. L.M., T.M.G., R.R., and D.J.I. analyzed and interpreted the data and wrote the manuscript. L.M and N.M generated yeast mimotope libraries. L.M. and T.M.G performed the experiments. B.C and I.S assisted with mimotope identification. D.M.M. mined single cell RNA-sequencing data. L.M., L.Z., and R.S. carried out amphiphile-mimotope synthesis. R.R. performed structural modeling. I.S. P.Y. and H.S. assisted with sample preparation. W.A., A.N., R.T., and A.C. assisted with animal experiments. A.R. performed lymph nodes immunofluorescence studies. B.G. performed ELISA analysis. S.K., R.M.M., S.A.G., L.P., G.G., S.J.S, N.F. and M.R. processed and provided the clinical data. E.B. and R.C assisted with ALK mimotope identification and verification.

