## Supplementary material for "Directed evolution-based discovery of ligands for in vivo restimulation of CAR-T cells": Materials and methods and supplemental figures

### Clinical trial data

*Pediatric B-cell acute lymphoblastic leukemia*

Patient data from two studies of tisagenlecleucel for the treatment of CD19-positive relapsed or refractory B-acute lymphocytic leukemia (B-ALL) in children and young adults, NCT01626495 [1] (N=60 infused) and NCT02906371 (n=70 infused) [2] were pooled (n=130). In both studies, patients received lymphodepleting chemotherapy with fludarabine and cyclophosphamide followed by a pre-infusion bone marrow aspirate or biopsy to assess tumor burden, and infusion of tisagenlecleucel over 1-2 days. Patients were categorized based on the pre-infusion bone marrow aspirate or biopsy as high (≥40%, n=40) or low tumor burden (<40%, n=90). Peripherally circulating CD19 CAR-T cells were quantified by quantitative polymerase chain reaction (qPCR). Peripherally circulating normal B cells were quantified by flow cytometry. Data analyses were performed using R. The study protocols were approved by the institutional review boards of the Children's Hospital of Philadelphia and the University of Pennsylvania. Patients or their guardians provided written informed consent.

*Adult B-cell acute lymphoblastic leukemia*

We conducted a retrospective analysis of CAR-T expansion in a cohort of adult B-ALL patients treated with the 4-1BB CART19 CTL019 (NCT02030847), now commercially known as tisagenlecleucel. Only patients with available pre-CART19 bone marrow involvement evaluation were included (n=29). Patients' characteristics were retrieved from the clinical trial database and electronic medical records as per an IRB-approved protocol. Tumor burden was defined as the percentage of bone marrow blast involvement before CAR-T infusion. Patients with more than 5% of blasts were defined as high tumor burden (n=23), while patients with ≤5% blasts were considered low tumor burden (n=6). Peak of CAR-T expansion was defined as the highest value of CAR19 copies/µg DNA after CAR-T infusion, which typically occurs between days 7 and 14. CAR-T cell expansion in the peripheral blood was evaluated using qPCR at multiple time points, as previously described[3,4]. A Student's t-test was used to compare CAR-T peak. Significance was set at 0.05.

*Adult B cell lymphoma*

We conducted a retrospective analysis of CAR-T expansion, according to tumor burden, in a cohort of B-cell Lymphoma patients treated with 4-1BB CART19 CTL019 (NCT02030834), now commercially known as tisagenlecleucel. Only patients with LDH levels available at CART19 infusion were included (n=37). Tumor burden was estimated per LDH serum levels at infusion day. Patients with LDH levels higher than 1.5-fold the upper normal limit were considered as high tumor burden (n=7). Peak of CAR-T expansion was defined as the highest value of CAR19 copies/µg DNA after CAR-T infusion, which typically occurs between days 7 and 14. CAR-T cell expansion in the peripheral blood was evaluated using qPCR at multiple time points, as previously described[3,4]. Patients' characteristics were retrieved from the clinical trial database and electronic medical records as per IRB-approved protocols. A t-test was used to compare CAR-T peak. Significance was set at 0.05.

### Cell line and animals

K562, Jurkat, NALM6 and 293 phoenix cells were obtained from ATCC. The NALM6-Luc cell line was a gift from Dr. Michael Birnbaum at MIT. The Eµ-Myc cell line was a gift from Dr. Michael Hemann at MIT. Wildtype female C57BL/6 mice ($CD45.2^+$), $CD45.1^+$ congenic mice, B6(Cg)-Tyrc-2J/J (C57BL/6J albino) and NSG mice were purchased from the Jackson Laboratories. All animal studies were carried out following an IACUC-approved protocol following local, state, and federal guidelines. The EBY100 yeast strain was a gift from lab of K. Dane Wittrup at MIT.

**Antibodies and recombinant proteins**

FMC63 IgG, his-tagged FMC63 scFv, 1D3 IgG, anti-ALK IgG (ALK123) were cloned into the gWiz vector and used for recombinant protein production using Expi293 cells. Recombinant proteins were then purified using FPLC (Akta pure 25). Anti-hCD19 (HIB19) and anti-HA antibody were purchased from Biolegend. The anti-Myc antibody (9B11) was purchased from Cell Signaling.

**Yeast mimotope library construction**

We followed previously established protocols[5,6] to construct our yeast surface display libraries starting with the parental pCT-Con2 vector, except the following modifications that were tailored for the mimotope library: 1) The mimotope library PCR was carried out with nested PCRs. 1st PCR primers: Mimotope Lib F - AACTAGCAAAGGCAGCCCCATAAACAC and Mimotope Lib R- GATTTTGTTACATCTACACTGTTGTTATCAGATCTCGAGCTATTAMNNMNNMNNMNNMNNMNNMNNMNNMNNMNNMNNGCTAGCCGACCCTCCGCC. 2nd PCR primers: Min Cmyc EP F – GGCTCTGGTGGAGGCGGTAGCGGAGGCGGAGGGTCGGCTAGC and Min Cmyc EP R -GATTTTGTTACATCTACACTGTTGTTATCAGATCTCGAGCTATTA. PCR products were verified on a 2.5% agarose gel. The nested PCR products were extracted using the conventional pheno-chloroform and ethanol precipitation method, pelleted and dissolved in 15 µl $H_2O$. The parental pCT-Con2 vector (Addgene # 41843) was digested with NheI, SalI and XhoI, purified using gel extraction followed by a clean-up step using PCR clean-up columns and eluted in 20 µl $H_2O$. For electroporation, 4 µg digested backbone was pre-mixed with the entire PCR product with the maximum volume less than 20µl. Then, pre-chilled yeast cells were added to the DNA mixture before transferring to an electroporation cuvette.

To generate mimotope library V2 with a fixed motif (RXCPWXCXXX), the mimotope Lib R primer was replaced with mimotope Lib V2 R -GATTTTGTTACATCTACACTGTTGTTATCAGATCTCGAGCTATTAMNNMNNMNNACAMNNCCACGGACAMNNACG GCTAGCCGACCCTCCGCCTC. To generate mimotope library V3 with extended flanking sequences ($X_{0\text{-}6}$RICPWNCKEL$X_{3\text{-}10}$), mimotope Lib R primer was replaced with:

mimotope Lib V3 R1 - GATTTTGTTACATCTACACTGTTGTTATCAGATCTCGAGCTATTA MNNMNNMNNAAGCTCCTTACAATTCCACGGACAAATACGMNNMNNMNN GCTAGCCGACCCTCCGCCTC, to generate library $X_3$RICPWNCKEL$X_3$.

mimotope Lib V3 R2 - GATTTTGTTACATCTACACTGTTGTTATCAGATCTCGAGCTATTA AAGCTCCTTACAATTCCACGGACAAATACGMNNMNNMNNMNNMNNMNN GCTAGCCGACCCTCCGCCTC, to generate library $X_6$RICPWNCKEL.

mimotope Lib V3 R3 - GATTTTGTTACATCTACACTGTTGTTATCAGATCTCGAGCTATTA MNNMNNMNNMNNMNNMNNAAGCTCCTTACAATTCCACGGACAAATACG GCTAGCCGACCCTCCGCCTC, to generate library XRICPWNCKEL$X_6$.

mimotope Lib V3 R4 - GATTTTGTTACATCTACACTGTTGTTATCAGATCTCGAGCTATTA MNNMNNMNNAAGCTCCTTACAATTCCACGGACAAATACGMNNMNNMNNMNNMNNMNN GCTAGCCGACCCTCCGCCTC, to generate library $X_6$RICPWNCKEL$X_3$.

mimotope Lib V3 R5 - GATTTTGTTACATCTACACTGTTGTTATCAGATCTCGAGCTATTA MNNMNNMNNMNNMNNMNNAAGCTCCTTACAATTCCACGGACAAATACGMNNMNNMNN GCTAGCCGACCCTCCGCCTC, to generate library $X_3$RICPWNCKEL$X_6$.

mimotope Lib V3 R6 - GATTTTGTTACATCTACACTGTTGTTATCAGATCTCGAGCTATTA MNNMNNMNNMNNMNNMNNMNNMNNMNNMNNAAGCTCCTTACAATTCCACGGACAAATACGGCTAGCCGACCCTCCGCCTC, to generate library RICPWNCKEL$X_{10}$.

mimotope Lib V3 R7 - GATTTTGTTACATCTACACTGTTGTTATCAGATCTCGAGCTATTA AAGCTCCTTACAATTCCACGGACAAATACGMNNMNNMNNMNNMNNMNNMNNMNNMNNMNNGCTAGCCGACCCTCCGCCTC, to generate library $X_{10}$RICPWNCKEL. A 10x diversity (definition of diversity: total number of mimotope variants within the yeast library) of all individual V3 libraries were mixed to create the final $X_{0\text{-}6}$RICPWNCKEL$X_{3\text{-}10}$ library.

### Yeast surface display screen

To prepare biotinylated recombinant protein, 200 µl of antibody or scFv at 1 mg/ml in PBS was mixed with 10% volume of 1M sodium bicarbonate to adjust pH to 7-8 followed by addition of EZ-Link sulfo-NHS-LC-Biotin (ThermoFisher) to reach a 5:1 biotin:protein molar ratio. The mixture was stirred at room temperature for 1 hour, followed by spin-column purification and subsequent quantification using nanodrop.

The yeast library was thawed, passaged and prepared as previously described[5,6]. To prepare for the screen, yeast cells were grown in SD-CAA until the OD600 reached 6-8. 30x diversity (or at least 100M) yeast cells were transferred into SG-CAA medium supplemented with 5% SD-CAA. Yeast were shaken at 20°C overnight to induce protein display on the cell surfaces. We noticed that 5% of SD-CAA improved mimotope display on the yeast cell surface, and usually >75% induction could be achieved.

The yeast library screen was performed as previously described[5,6] with the following modifications: 1) 15x diversity was used per sort (7.5e9 for the mimotope library). Yeast was spun down at 3500RPM for 5 minutes, washed in cold 1x PBSA (1x PBS + 0.1% BSA), then aliquoted into 2 x 2 mL tubes (3.75e9 per tube). Yeast was then spun at 12,000xg for 1 minute and resuspended in 1mL 1x PBSA. 2) For positive-sort beads (Dynabeads™ Biotin Binder [ThermoFisher] pre-coated with 6.7-33pmoles of biotinylated IgG or scFv as previous described[6]), non-bound yeast was gently removed, and the remaining beads were washed to remove unbound or weakly bound yeast cells with 1ml 1x PBSA by inverting repeatedly. Supernatant was removed, then yeast were resuspended in 1 mL SD-CAA, and counted to determine diversity. 3) 10x diversity of 1$^{st}$ round positive-sorted yeast was pelleted at 12,000xg for 1 minute. Yeast cells were inoculated into premade 5 ml SG-CAA with 5% SD-CAA, then induced at 20°C overnight (at least 8 hr is needed for reasonable induction). At least 20x diversity was pelleted and washed once with 1x PBSA, then resuspended in 1ml 1x PBSA. 4) For the 2$^{nd}$ round of magnetic enrichment, the first bare bead sort and two subsequent negative sorts using beads coated with isotype control IgG were carried out at 0.5 beads per yeast cell for >=2 hours at 4°C. This step was critical to remove background and non-specific binders. The final positive sort using FMC63$_{IgG}$-coated beads was carried out following the standard enrichment procedure but with longer washes (30 sec) to improve stringency. Enriched yeast cells were counted under microscopy to determine the diversity. At this point, the diversity should have been massively reduced. 5) 30x of enriched and induced yeast cells from the 2$^{nd}$ round of positive sorting were pelleted and stained with 30 µl 5 µM control IgG or FMC63 IgG at 4°C for 30 minutes. Yeast was washed twice with 1 ml of 1x PBSA to remove residual antibody. When a plate was used for staining, yeast was washed 3-4 times with 200 µl. The pellet was stained in 50 µl of 1:100 dilution PE-Streptavidin and BV421-HA for 20 minutes on ice. Yeast was washed twice with 1 ml of 1x PBSA prior to flow cytometry sorting (BD FACS Aria), with adjustments made as previously described for using a plate when needed. The top 0.5-1% of the major population based on FMC63 IgG binding was sorted. Usually following this modified protocol, a clearly distinct yeast population could be observed during flow cytometry analysis. 6) FMC63 IgG was used at 0.5µM for the subsequent flow cytometry-based sort. We alternated between streptavidin and anti-biotin antibodies when staining yeast populations for flow cytometry to avoid selecting streptavidin binders. 7) For kinetic sorting, 10x of the library V3 was stained with 50nM biotinylated FMC63 scFv for 30min, washed 2x with 1x PBS and then incubated with 500 nM of non-modified FMC63 IgG for 1 hour or overnight.

### Plasmid extraction from yeast cells and sanger sequencing

The Zymo yeast DNA extraction kit was used with a modified protocol. Briefly, 50 million yeast cells were pelleted and resuspended in 200 µl solution I. 3 µl of zymolyase was added, then the mixture was vortexed and incubated at 37°C for 1 hour. 200 µl of solution 2 was added, then mixed gently by flipping tubes upside down a few times, and leaving at room temperature for 5 min. 400 µl of solution 3 was added, then mixed well by shaking. The mixture was spun at top speed for 5 minutes. Supernatant was transferred to a new 1.5 ml EP tube and spun at top speed for 5 minutes to remove all white clumps. The clear lysate was transferred onto a spin column from a regular miniprep kit instead of the column provided in the Zymo yeast DNA extraction kit, then spun at 11,000g for 30 seconds. Columns were washed with 600 µl of wash buffer from the miniprep kit, spun again at 11,000g for 1 minute to dry the membrane, and then eluted with 50 µl of $H_2O$. Eluted DNA was cleaned using a PCR clean-up kit (Invitrogen) following the manufacturer's protocol. DNA was eluted in 20 µl $H_2O$ and quantified using nanodrop. 5 µl of clean DNA was transformed into 50 µl of DH5α commercial competent cells (NEB). The whole transformation was spread onto a 10 cm ampicillin agar plate. Individual bacteria colonies were picked and inoculated onto a 96-well agar plate and grown overnight. The entire plate was sent for miniprep and sanger sequencing.

**ELISA**

Mouse and human ELISA assays were performed following the manufacturer's protocol (Mouse and human IFN-g and TNF-Duo set, R&D systems). For ELISA monitoring of FMC63 IgG binding to mimotopes, mimotope was dissolved in 1x PBS and coated onto a 96-well maxisorp plate at 4 µg/ml, 50 µl per well, sealed and kept at 25°C overnight. The next day, the plate was washed 4x followed by 1hr blocking (1% BSA in 1x PBS) at 25°C. After washing, FMC63 IgG was diluted to desired concentration in the blocking buffer and added to each well at 50 µl per well. After 1hr incubation at 25°C, plates were washed again, substrate was added and the remainder of the assay was performed as previously described[7]. Anti-mimotope antibody response was determined by ELISA. Maxisorb plates (Thermo Fisher) were coated with 4 µg/ml of mimotope in PBS overnight at room temperature, washed 3X with 1x wash buffer (PBS containing 0.1% Tween-20(v/v), then blocked with blocking buffer (1% BSA in PBS) at 25°C for 1 hr. 50 µl of serum diluted 1:100 was transferred into each well, covered, and incubated for 1 hr at 25°C. Plates were then washed 3X, and 50µl per well of goat anti-mouse IgG-HRP (BioRad, 1:5000) was added, diluted in blocking buffer. Plates were incubated for 1 hr at 25°C, washed 3X, and 50µl per well of TMB Chromogen Substrate Solution (Thermo Fisher) was added. Plates were kept at 25°C in the dark. After about 20 min, the reaction was terminated with 50 µl of 2N H2SO4. Absorbances were read at 450 nm on a Tecan Spark plate reader (Tecan Life Sciences), with the subtraction of background reading at 540 nM. Mouse serum was collected on day 21.

**Construction of murine and human CARs**

Murine CAR-expressing constructs were generated by fusing geneblock fragments (custom ordered from IDT) into an MSCV retroviral vector. The hybrid FMC63-mCAR sequence is composed of a mouse CD8 signal peptide, FMC63 scFv, mouse CD8a hinge and transmembrane domain, CD28 costimulatory domain and CD3ζ intracellular domain as described in our previous work[7]. The tandem αFITC-FMC63-mCAR was constructed as described in our previous work [7] by fusing an αFITC scFV (clone 4m5.3) to the N-terminus of the FMC63-mCAR (CD28) via a (G4S)4 linker with the mouse CD8 signal peptide at the N-terminus of the αFITC scFV, and a Myc tag was inserted immediately after the FMC63 scFv and before the mouse CD8 hinge to facilitate the detection of CAR expression on T cell surface. The human CAR-expressing constructs were generated by fusing geneblock fragments (custom ordered from IDT) into a lentiviral vector containing the EF1a promoter. The CD19-CAR is composed of a human CD8 signal peptide,

FMC63 scFv (VLVH), human CD8 hinge and transmembrane domain, CD28 costimulatory domain and CD3ζ intracellular domain as described previously[7]. To facilitate CAR detection by flow cytometry, a Myc tag was inserted at the N-terminus of FMC63 scFv immediately following the signal peptide in both constructs.

**Virus production**

For optimal retrovirus production, 293 phoenix cells were cultured till 80% confluence, then split at 1:2 for further expansion. 24 hr later, $5.6 \times 10^6$ cells were seeded in a 10 cm dish and cultured for 16 hr till the confluency reached 70%. 30 min – 1 hr before transfection, each 10 cm dish was replenished with 10 ml pre-warmed medium. Transfection was carried out using the calcium phosphate method following the manufacturer's protocol (Clontech). Briefly, for each transfection, 18μg of plasmid (16.2 μg of CAR plasmid plus 1.8 μg of Eco packaging plasmid) was added to 610 ml of dd$H_2O$, followed by addition of 87ml of 2 M $CaCl_2$. 700ml of 2x HBS was then added in a dropwise manner with gentle vortexing. After a 10 min incubation at 25°C, the transfection mixture was gently added to phoenix cells. After 30 min incubation at 37°C, the plate was checked for the formation of fine particles, as a sign of successful transfection. The next day, old medium was removed and replenished with 8 ml of pre-warmed medium without disturbing the cells. Virus-containing supernatant was collected 36 hr later and passed through a 0.45 um filter to remove cell debris, designated as the "24hr" batch. Dishes were refilled with 10 ml of fresh medium and cultured for another 24 hr to collect viruses again, designated as the "48hr" batch, this process can be repeated for another two days to collect a "72hr" batch and "96hr" batch. All virus supernatant was aliquoted and stored at -80°C. Virus transduction rate was evaluated in a 12-well format by mixing 0.5 million activated T cells with 0.5ml of viruses from each batch. Plate coating, spin infection and flow cytometry analysis of CAR expression were carried out as described below. In the majority of experiments, the "48hr" and "72hr" batches yielded viruses that transduced T cells at 90-95% efficiency, the "24hr" and "96hr" batch viruses led to >80% transduction. Only viruses with >80% transduction rate were used for animal studies. Lentivirus was produced as previously described[8].

**Primary mouse T cell isolation and CAR-T cell production**

For T cell activation, 6-well plates were pre-coated with 5 ml of anti-CD3 (0.5 μg/ml, Clone: 2C11) and anti-CD28 (5 μg/ml, Clone: 37.51) per well at 4°C for 18 hr. $CD8^+$ T cells were isolated using a negative selection kit (Stem Cell Technology), and seeded onto pre-coated 6-well plates at 5 $\times 10^6$ cells/well in 5 ml of complete medium (RPMI + penicillin/streptomycin + 10% FBS + 1x NEAA + 1x Sodium pyruvate + 1x 2-mercaptoethanol + 1x ITS [Insulin-Transferrin-Selenium, ThermoFisher]). Cells were cultured at 37°C for 48 hr without disturbance. 24 hr before transduction, non-TC treated plates were coated with 15 μg/ml of retronectin (Clontech). On day 2, cells were collected, counted and resuspended at $2 \times 10^6$ cells/ml in complete medium supplemented with 20 μg/ml of polybrene and 40 IU/mL of mIL-2. Retronectin-coated plates were blocked with 0.05% FBS containing PBS for 30 min before use. 1 ml of virus supernatant was first added into each well of the blocked retronectin plate, then 1 mL of the above cell suspension was added and mixed well by gentle shaking to reach the working concentration of polybrene at 10 μg/ml and mIL-2 at 20 IU/ml. Spin infection was carried out at 2000x*g* for 120 min at 32°C. Plates were then carefully transferred to an incubator and maintained overnight. On day 3, plates were briefly centrifuged at 1,000x*g* for 1 min, and virus-containing supernatants were carefully removed. 3 mL of fresh complete medium containing 20 IU/ml of mIL-2 were then added into each well. Cells were passaged 1:2 every 12 hr with fresh complete medium containing 20 IU/mL of mIL-2. Transduction efficiency was evaluated by surface staining of a c-Myc tag included in the CAR construct[7] using an anti-Myc antibody (Cell signaling, Clone:9B11) ~30 hr after transduction. If needed, on day 3, after flow cytometry analysis of virus transduction, CAR-T cells could be frozen down and stored for assays at a later time. For in vivo experiments, CAR-T cells were used on day 4. For *in vitro* experiments, CAR-T cells were cultured until day 5.

**Primary human T cell isolation and CAR-T cell production**

Buffy coats were obtained from anonymous healthy donors (Research Blood Components, LLC, Boston, MA, USA). Total peripheral blood mononuclear cells (PBMCs) were isolated by Ficoll-Paque PLUS gradient separation. $CD8^+$T cells were isolated directly using the EasySep Human $CD8^+$ T cell isolation kit (Stemcell). For experiments completed at the Children's Hospital of Philadelphia (CHOP), $CD3^+$ T cells were isolated by negative selection using RosettaSep Kits from STEMCELL Technologies and obtained from Human Immunology Core at the Perelman School of Medicine at the University of Pennsylvania. T cells were activated with Human T-Activator CD3/CD28 Dynabeads (ThermoFisher) at a bead-to-cell ratio of 3:1 in complete medium supplemented with 30 IU/mL recombinant human IL-2 (PeproTech). After two days of activation, T cells were transduced with lentiviral supernatants as described above for murine T cells, and transduction efficiencies were determined by flow cytometry 2 days later. If transduction efficiency was less than 50%, $CAR^+$ T cells were enriched by staining total expanded T cells with a PE-conjugated anti-Myc antibody followed by staining with anti-PE microbeads and magnetic selection for $CAR^+$ T cells. Enriched CAR-T cells were continuously expanded till day 7 for adoptive transfer to NSG mice.

**Mimotope, amphiphile-mimotope production and vaccination**

Various mimotopes were custom synthesized (GenScript). The mimotope with a thioacetal bond was synthesized according to a previously published protocol[9] and verified by MALDI. Amph-mimotope molecules were produced and purified as previously described with modifications as indicated below[7]. Briefly, mimotope peptides (with an N-terminal Azido lysine) were dissolved in $H_2O$ at 10 mg/mL and mixed with 1.1 molar equivalent of DSPE-PEG2k-DBCO (Avanti). The mixture was agitated at 25°C for 24 hr. Conjugation efficiency was analyzed using a C4 column on HPLC (Schimadzu). Typically, the conjugation efficiency reached >95%. When the conjugation efficiency was low, unconjugated peptide was removed using HPLC and the conjugates were collected. The resulting products were lyophilized, re-dissolved in PBS, quantified using nanodrop and stored at -20°C. DSPE-PEG-FITC was purchased from Avanti. For vaccination, unless otherwise stated, mice received weekly s.c. injection of 10 µg peptide equivalent of amph-mimotope mixed with 25 µg of Cyclic-di-GMP (CDG, Invivogen) in 100 µl 1x PBS, administered 50 µl to each side at the tail base.

***In vitro* and *in vivo* amph-mimotope labeling**

For *in vitro* labeling, target cells were pelleted at 1,000x*g* for 3min and washed with PBS twice to remove residual protein. The cell pellet was then resuspended at $1x10^6$ cells/ml in PBS containing amph-mimotope molecules at the indicated concentrations and incubated at 37°C for 30 min. The labeling reaction was stopped by pelleting cells and washing with PBS twice. For *in vivo* labeling, mice received 10µg peptide equivalent of amph-mimotope with or without CDG as described above. 24 hr later, Inguinal LNs were extracted and dissociated into single cell suspension for flow cytometry staining for macrophages ($MHCII^+CD11b^+CD11c^-F4/80^+$), cDC1($MHCII^+$ $CD11c^+CD11b^{low/-}CD24^+$) and cDC2 ($MHCII^+CD11c^+CD11b^+CD24^{low/-}$) as previously described[10]. To detect amph-mimotope decoration of various lymph node cell populations, 100nM of biotinylated FMC63 IgG was included in the antibody cocktail followed by secondary staining with AlexaFluor 647-streptavidin.

**CAR-T functionality assay**

The functionality of CAR-T cells was assessed by co-coculturing with target cells *in vitro*. 96-well U bottom plates were used. Unless otherwise stated, $1x10^5$ sorted CAR-T cells or unsorted CAR-

T cells (if >70% CAR+ T cells) possessing an equivalent number of $CAR^+$ T cells were mixed with $1x10^4$ target cells in a total volume of 200μl complete medium containing 20IU/ml of mIL-2 (for mouse hybrid CAR-T) or 30IU/ml of hIL-2 (for human CAR-T). After 6 hr co-culture, cells were pelleted at 2,000x*g* for 5 min, and supernatants were harvested for IFN-γ ELISA.

**Modeling mimotope interaction with FMC63**

Both F12 and F12-A1 were modeled as multimers using Alphafold with default settings[11]. Alphafold generated 25 structures that were relaxed using the AMBER force field option, then ranked internally. All 25 structures were clustered with Rosetta[12] using an RMSD cut-off of 2.0Å to better understand which conformations were favored during modeling. For both F12 and F12-A1, the top ranked model resided in the cluster with the most members, and the most favorable energy when scored using the Rosetta score function[13,14]. The top ranked structure was minimized using the Rosetta FastRelax[15] protocol with deviations up to 3.0Å allowed, to eliminate any clashes between sidechains and achieve a more favorable conformation when scored using Ref2015. FastRelax produced 100 constructs, of which the construct with the most favorable energy was designated as the candidate structure. PDB 7URV was similarly minimized, with deviations up to 0.5Å allowed, to ensure coordinates are constrained to the crystal structure. All interfaces were analyzed using the PDBePISA webserver[16].

***In vivo* tracking of CAR-T proliferation and response to amph-mimotope stimulation in syngeneic mouse**

To monitor short-term amph-mimotope stimulation of CAR-T cells *in vivo*, $2x10^6$ murine hybrid FMC63-mCAR-T cells and untransduced T cells were mixed at a 1:1 ratio and labeled with 2.5 μM CTV and i.v. infused into recipient C57BL/6 mice. 16 hr later, 100 μl of amph-mimotope vaccine (10 nmol of amph-mimotope mixed with 25μg CDG in 100μl PBS) was s.c. injected into recipient mice, with 50 μl on each side of the tail base. After an additional 48 hr, mice were euthanized and inguinal LNs were excised for flow cytometry analysis. Live $CD3^+CD8^+CTV^+$ cells were gated as donor cells, and staining for the Myc tag on the CAR was used to distinguish CAR-T cells from non-CAR-T cells. For long-term monitoring of hybrid FMC63-mCAR-T cell expansion in response to amph-mimotope vaccination, recipient CD45.2 mice received sublethal lymphodepletion (500cGy gamma irradiation) on day -1 followed by i.v. infusion of $1x10^6$ CD45.1 FMC63-mCAR-T cells on day 0. Two weekly doses of amph-mimotope vaccines were given on day 1 and day 7, peripheral blood was sampled on day 4, day 7 and then every week thereafter for enumeration of CAR-T cells.

**Preparation of monocyte-derived DCs and amph-mimotope labeling**

Peripheral blood mononuclear cells were isolated from a buffy coat using Ficoll-Paque density gradient. $CD14^+$ monocytes were then purified using the pan monocyte isolation kit (Miltenyi BioTec) or isolated by negative selection using RosettaSep Kits from STEMCELL Technologies and obtained from Human Immunology Core at the Perelman School of Medicine at the University of Pennsylvania. The characterization and differentiation of monocyte to immature DCs were performed as previously described[17]. DC maturation was carried out using LPS and IFN-γ as reported previously[18]. To label mature DC for *in vivo* vaccination or *in vitro* killing assay, mature DCs were washed with 1x PBS twice and then stained with 500 nM of amph-mimotope.

**Immunocompetent B-ALL/Lymphoma mouse model**

To develop an immunocompetent mouse model for CAR-T cell therapy, we utilized a tumor cell line derived from an Eμ-Myc transgenic mouse, which develops B-ALL and Burkitt-like B cell lymphoma-like malignancy[19,20]. Eμ-Myc cells were modified to express human CD19 (hCD19) using a retroviral vector. Purification of transduced cells was performed using anti-PE MicroBeads (Miltenyi Biotec) and PE-conjugated anti-human CD19 antibody. To enable monitoring of disease progression, hCD19$^+$ Eμ-myc cells were transduced with retroviral vector encoding mCherry and firefly luciferase. Cells were cultured in a medium composed of a 50:50 mix of IMDM with L-glutamine and 25 mM HEPES (Gibco) and DMEM with L-glutamine and sodium pyruvate (Corning), supplemented with 10% FBS and 2-mercaptoethanol to a final concentration of 0.05 mM (Gibco).

All animal work in this model was conducted under with the CHOP Department of Veterinary Services (DVR) under an animal protocol approved by the Institutional Animal Care and Use Committee at CHOP in accordance with federal, state, and local guidelines. B cell lymphomas were established by i.v. injection of 0.5 × $10^6$ Eμ-Myc cells in C57BL/6J or B6(Cg)-Tyrc-2J/J (C57BL/6J albino, Jackson Laboratory) mice after sublethal irradiation (500 cGy X-ray irradiation). On day 4, animals received either mock treatment (untransduced CD45.1$^+$ T cells), FMC63-mCAR-T, or αFITC-FMC63-mCAR-T cells (2x$10^6$) intravenously, followed by three rounds of weekly s.c. immunization on both sides of the tail base with amph-mimotope vaccine (10 μg of amph-mimotope mixed with 25 μg CDG in 100 μl PBS, 50ul each side) or amph-FITC vaccine (10 nM of amph-FITC mixed with 25 μg CDG in 100 μl PBS), respectively. Disease progression was subsequently monitored every 2-3 days using an IVIS Spectrum fluorescence/bioluminescence imaging system (PerkinElmer) with intraperitoneal administration of 150mg $kg^{-1}$ d-luciferin $K^+$ salt (PerkinElmer catalog # 122799). Total photon counts or radiance was quantified using Living image 4.5. Mice were monitored daily and euthanized at morbidity or as recommended by the veterinarian. The toxicity of therapy was monitored using a toxicity score as described before[21]: A toxicity score of 0 corresponded to active, well-groomed animals, with well-kept hair coat and invisible spine. A score of 1 was assigned to mice with signs of hypomotility, tousled hair coat and a partially visible spine. A score of 2 was given in case of somnolence, rough, dull or soiled hair coat, hunched back with visible spine. Peripheral blood was collected on days 11 and 18. CAR-T expansion and immunophenotyping of CAR-T cells was carried out by flow cytometry. Red blood cells were lysed in ACK Lysis Buffer (Thermo Fisher) before flow cytometry staining followed by a surface staining for CD45.1 (BV421, clone: A20), CD62L (PE-Cy7, clone: MEL-14), CD44 (BV711, clone: IM7). The number of cells was determined using CountBright Plus Absolute Counting Beads (Thermo Fisher). The serum level of IFN-γ, TNF-α, IL-10, IL-6, and IL-2 was determined using a bead-based multiplex assay panel, LEGENDplex Mouse Th1 panel (Biolegend), according to the manufacturer's protocol.

**Intracelullar cytokine staining (ICS)**

Peripheral blood (PB) was collected from mice receiving CAR-T or CAR-T plus booster vaccines at day 6 post-vaccination. 100 μl PB was processed in ACK lysis buffer, PMBCs resuspended in 100 μl RPMI1640 medium with 10% FBS and 2X Golgi plug (Biolegend). $10^5$ Eμ-Myc hCD19$^+$ target cells were resuspended in RPMI1640 medium with 10% FBS. 100 μl of target cells was mixed with 100μl of PBMCs, transferred to 96-well flat-bottom plates and cultured at 37°C for 6 hr. As a positive control, extra PMBCs from mice receiving CAR-T were combined and cultured with both 1X Golgi plug and cell stimulation cocktail for 6 hr. Cells were then resuspended and transferred to 96-well V-bottom plate for downstream processing. Cells were pelleted and washed once with PBS, and stained with live/dead aqua for 15 min in the dark at 25°C. Cells were pelleted again, surface stained for CD45.1 (PerCP, clone:A20) for 20 min on ice followed by 1 wash with flow cytometry buffer. Cells were resuspended in 75 μl of BD Fix/Perm and kept at 4°C for 15 min, then washed once by direct filling with 200 μl 1x Perm/Wash (Thermo Fisher). The pellet was resuspended in 50 μl of cytokine antibody cocktail (IFN-γ (BV421, clone:XMG1.2) at 1:100, TNF-α (PE-Cy7, clone:MP6-XT22) at 1:100) pre-diluted in 1x Perm/Wash buffer, 30 min on ice, then washed once with 1x Perm/Wash buffer and resuspended in 1x flow cytometry buffer for analysis immediately or kept at 4°C for analysis on a BD Fortessa X-20 flow cytometer the next day.

For intracellular staining from CD19 CAR-T cell-treated NSG mice, splenocytes were resuspended in 200 µl RPMI1640 medium with 10% FBS, Golgi plug (Biolegend), and eBioscience™ Cell Stimulation Cocktail (ThermoFisher), transferred 96-well flat-bottom plates and cultured at 37°C for 6 hr. Cells were then transferred to 96-well V-bottom plate for downstream processing. Cells were pelleted and washed once with PBS then stained with live/dead aqua for 15 min in the dark at 25°C. Cells were pelleted again, surface stained for CD3 (PerCP-eFluor710, clone: OKT3) and Myc-tag (AlexaFluor 647, clone: 9B11) for 20 min on ice followed by 1 wash with flow cytometry buffer. Cells were resuspended in 75 µl of BD Fix/Perm and kept at 4°C for 15 min, then washed once by direct filling with 200 µl 1x Perm/Wash (Thermo Fisher). The pellet was resuspended in 50 µl of cytokine antibody cocktail (IFN-γ (BV421, clone: 4S.B3) at 1:50, TNF-α (BV605, clone: Mab11) at 1:50) pre-diluted in 1x Perm/Wash buffer, 30 min on ice, then washed once with 1x Perm/Wash buffer and resuspended in 1x flow cytometry buffer for analysis immediately or kept at 4°C for analysis on a BD Fortessa X-20 flow cytometer the next day.

**Lymph node analysis**

For lymph node tissue section imaging, C57BL/6 mice were lymphodepleted (L.D.) and injected with $0.5 \times 10^6$ $hCD19^+$ Eµ-Myc cells. On day 7, mice were adoptively transferred with $2 \times 10^6$ FMC63-mCAR-T cells or control T cells, then vaccinated 1 day later with 10µg amph-F12-A1 (Vax). Inguinal (draining) lymph nodes were collected on day 15. Lymph nodes were flash frozen in tissue cutting medium and sectioned into 10 micron-thick sections on a Leica Cryostat and stored at -80. The sections were fixed in 10% formalin and permeabilized and blocked with Perm/Block solution containing 1% bovine serum albumin and 0.01% Triton-X, then washed in 1X PBS. The sections were then treated with Fc blocker (Innovex) and stained with antibody solution (1:100 anti-CD3e AF488 (Biolegend 100321), 1:100 anti-B220 AF594 (Biolegend 103254), and 1:75 anti-CD11c AF647 (BD Biosciences 565587) diluted in Perm/Block buffer in a humidity chamber for 1.5 hours. The sections were washed 3 times in PBS, then stained with 300 nM DAPI solution for 5 minutes. The slides were washed once more and mounted with Pro-long Diamond anti-fade solution. The tissue sections were imaged on a Leica Sp8 Laser Scanning Microscope with a 25x objective. Laser power was kept consistent across groups. Shown are representative images of lymph node sections, with magenta representing B cells, green representing T cells, and white representing CD11c DCs. Scale bars represent 200 microns.

**Human B-ALL mouse model and therapeutic studies**

All animal work was conducted under an MIT Division of Comparative Medicine Institute animal care or CHOP Department of Veterinary Services (DVR), used a committee-approved animal protocol by the Committee of Animal Care at MIT or Institutional Animal Care and Use Committee at CHOP and used a committee-approved protocol in accordance with federal, state, and local guidelines. 8-12 week-old NOD.Cg-$Prkdc^{scid}IL2rg^{tm1Wjl}$/SzJ (NSG) mice (Jackson Laboratory) were injected with $0.5 \times 10^6$ NALM6-Luc cells and randomly assigned into each treatment group. On day 4, leukemia-bearing NSG mice received either mock treatment or a suboptimal dose of CD19 CAR-T cells ($2 \times 10^5$ for **Fig.7**, or $5 \times 10^5$ for **Extended Data Fig.10**.) intravenously. 24 hours later, one group of CAR-T-treated mice also received amph-mimotope decorated MoDCs (DC-mVax) every four days. Leukemia progression was subsequently monitored every 4-7 days using a Xenogen IVIS fluorescence/bioluminescence imaging system (PerkinElmer) with intraperitoneal administration of 150mg $kg^{-1}$ d-luciferin $K^+$ salt (PerkinElmer 122799). Total photon counts or radiance were quantified using Living image 4.5. Mice were monitored daily and euthanized at morbidity, >20% weight loss, severe graft versus host disease, or as recommended by the veterinarian. To monitor CAR-T cell expansion and persistence, peripheral blood was collected retro-orbitally, 50µl from each mouse was used for each flow cytometry analysis. Red blood cells were lysed in ACK Lysis Buffer (ThermoFisher) prior to flow cytometry staining and the total number of PBMCs per microliter blood in each sample was estimated by cell counting under a microscope or using CountBright Plus Absolute Counting Beads (Thermo Fisher). Spleens were collected from mice receiving CAR-T or CAR-T plus DC-mVax on day 19. Splenocytes were stained with live/dead aqua, followed by a surface staining for CD3 (PerCP-eFluor647, clone: OKT3), CD4 (PE-Cy7, clone: RPA-T4), Myc tag (AlexaFluor 647, clone: 9B11), CD45RA (AlexaFluor 488, clone: HI100), CCR7 (PE, clone: G043H7) or stained intracellularly for cytokines as described above

**Data mining and bioinformatic analysis**

Single-cell RNA-sequencing data from bone marrow of human patients with acute lymphoblastic leukemia was obtained from GSE134759. Cells with under 500 unique genes detected were filtered, and the data was normalized by library size using the "NormalizeData" function in Seurat. Variable features were selected using the "FindVariableFeatures" function, followed by data scaling with the "ScaleData" function and principal components analysis using "RunPCA". The data was processed for batch correction using Harmony. Clusters were defined using the "FindClusters" function, and two-dimensional embeddings were generated using uniform manifold approximation and projection (UMAP). Differential gene expression analysis was performed using the "FindMarkers" function. Likely-malignant B cells were identified as two clusters of B cells that were primarily detected in samples with a diagnosis of "Diagnosis" or "Relapse". A score for costimulatory marker expression was computed using the "AddModuleScore" function in Seurat with the following list of genes: *CD80, CD86, TNFSF9, ICOSLG, TNFSF4, TNFSF18, TNFSF14.*

**Statistics, selection of animals and justification of sample size**

Statistical analyses were performed using GraphPad Prism 8. Animal survival was analyzed using log-rank (Mantel-Cox) test. All pair-wise comparisons were analyzed by student's t-test. Multi-group comparisons were carried out using a one-way ANOVA with Tukey's multiple comparisons test. Experiments that involved repeated measures over a time course, such as the total flux, were analyzed using a RM (repeated measures) two-way ANOVA based on a general linear model (GLM). The RM design included factors for time, treatment and their interaction. Tukey's multiple comparisons test was carried out for the main treatment effect. P-values are adjusted to account for multiple comparisons in both one-way ANOVA, and RM two-way ANOVA. We determined the size of samples for experiments involving either quantitative or qualitative data as previously reported[22]. Based on our previous experience with the animal models and as reported by others[19,23,24], we consider the therapy as significant if it increases the survival of animals up to 100% within 4 weeks, and >=5 animals per group is necessary to achieve this goal with 95% confidence interval and at 80% power.

### EXTENDED DATA FIGURES

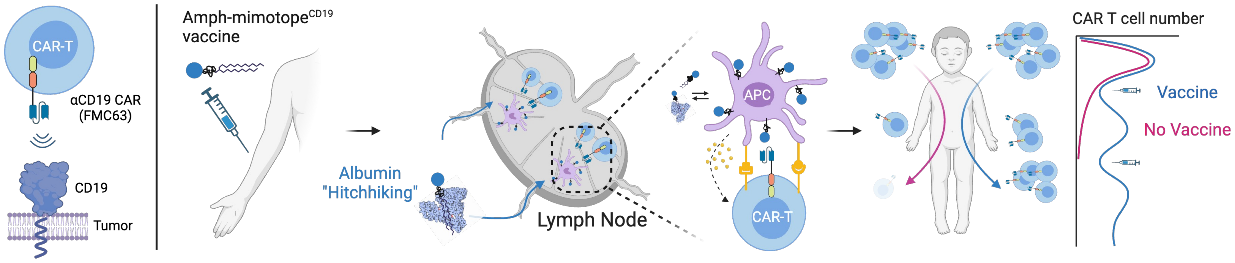


**Figure 1: The concept of designing an amph-mimotope vaccine for CD19 CAR-T cell therapy.**

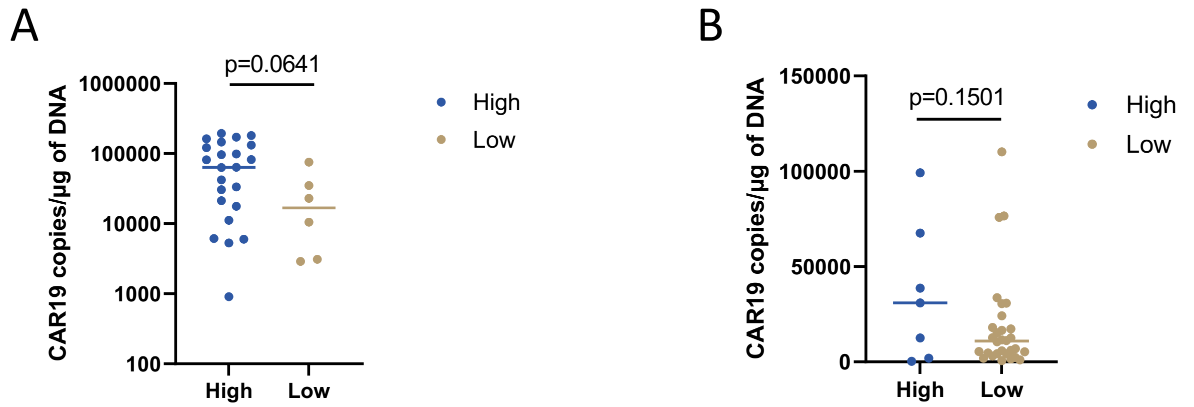


**Extended Data Figure 2. CD19 CAR-T cell expansion in Adult B-ALL and B-cell lymphoma patients with initial high or low tumor burden.** (**A**) The impact of high tumor burden on CAR-T expansion in a cohort of relapsed/refractory ALL patients (n=29) treated with 4-1BB CAR-T19 (CTL019; NCT02030847). High tumor burden was defined as bone marrow involvement before CART19 higher than 5%. The majority of patients (23/29, 79.3%) had a high disease burden before treatment. High tumor burden correlated with higher peak of expansion (CAR19 copies/µg of DNA: high tumor burden: 77,109 vs. low tumor burden: 25,064; p=0.0641;). (**B**) The impact of high tumor burden on CAR-T expansion in a cohort of relapsed/refractory B-NHL patients treated with 4-1BB CAR-T19 CTL019 within a clinical trial (n=37; NCT02030834). High tumor burden was defined as serum LDH higher than 1.5 folds the upper normal limit at infusion. In this cohort, we observed a trend toward higher CAR-T19 expansion in patients with high disease burden (n=7, 18.9%) than in patients with low tumor burden (n=30, 81.1%) (CAR19 copies/µg of DNA 35,857 vs 18,718 respectively; p=0.1501).

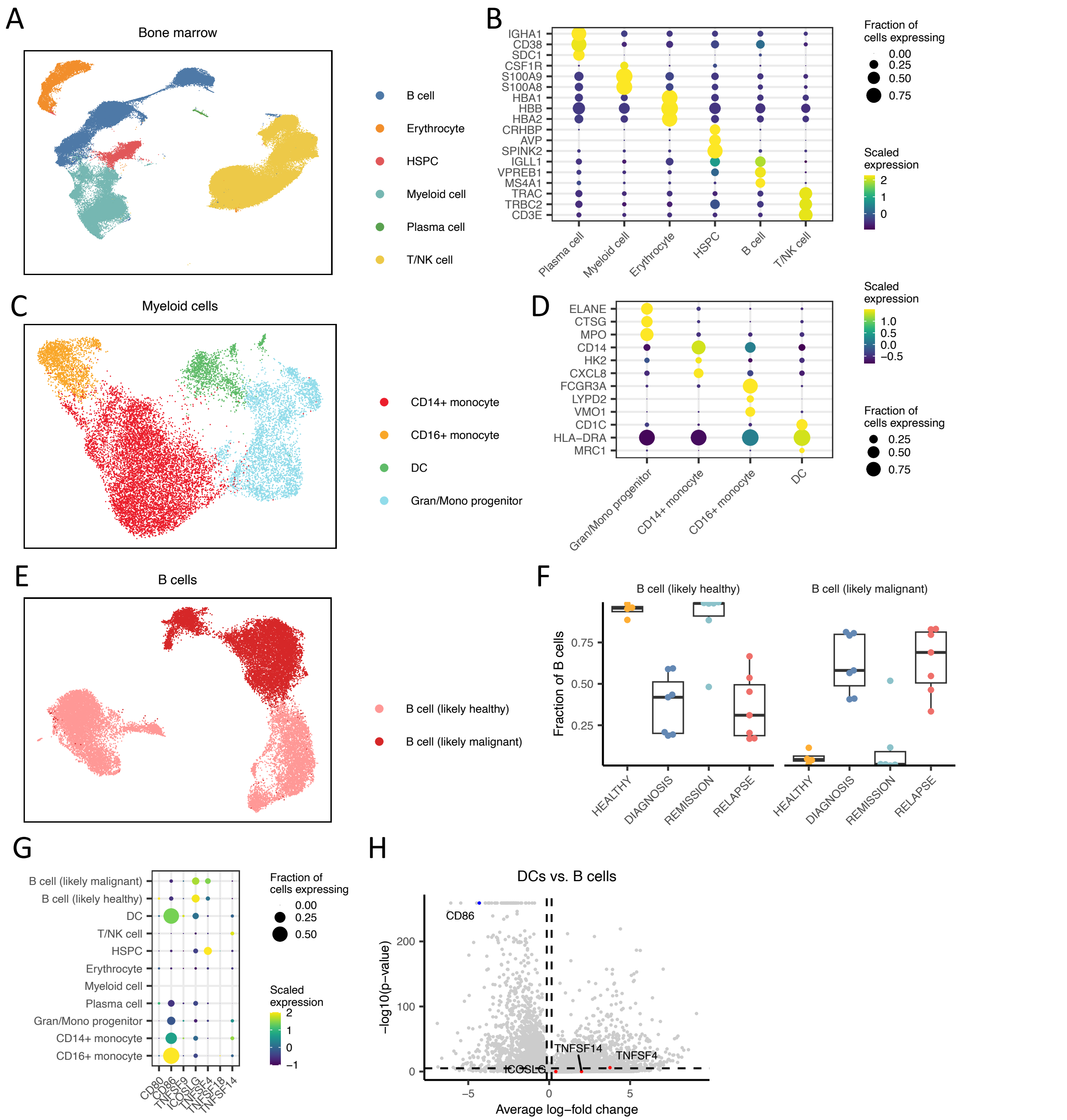


**Extended Data Figure 3. Single-cell RNA-seq analysis of immune cells from B-ALL patients.** (**A**) UMAP of single cells isolated from the bone marrow of human B-ALL patients, colored by cell phenotype. (**B**) Dot plot showing the scaled expression and fraction of cells expressing marker genes for each cell phenotype. (**C**) UMAP of myeloid cells, colored by cell phenotype. (**D**) Dot plot showing the scaled expression and fraction of cells expressing marker genes associated with each myeloid cell phenotype. (**E**) UMAP of B cells, with likely-healthy and likely-malignant cells distinguished. (**F**) Frequency of likely-healthy and likely-malignant B cells in samples with each diagnosis. (**G**) Dot plot showing the scaled expression and fraction of cells expressing each costimulatory receptor. (**H**) Volcano plot of genes differentially expressed between DCs and B cells. P-values are calculated with a two-sided Wilcoxon rank-sum test and are adjusted using Bonferroni correction

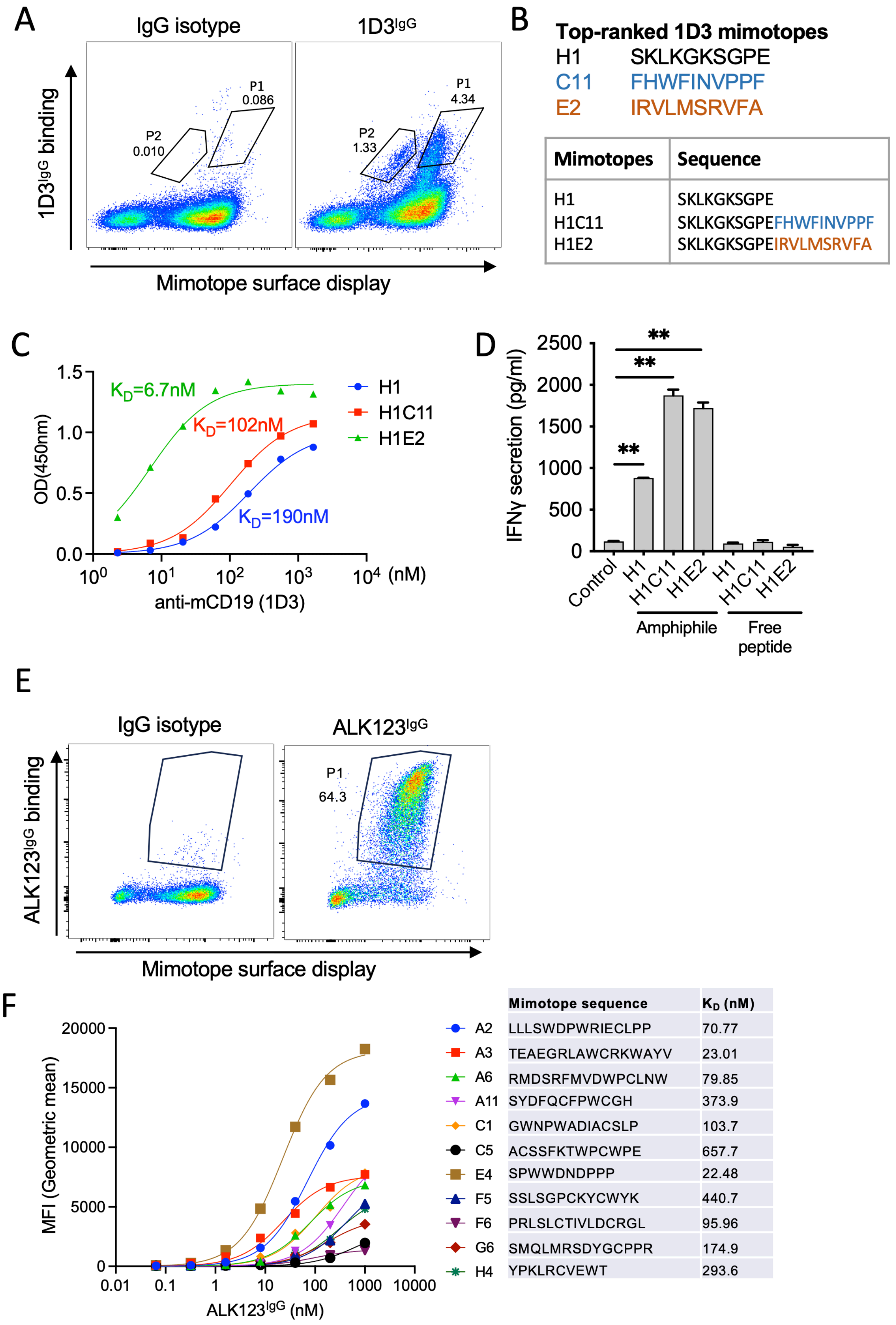


| Mimotopes | Sequence |
|---|---|
| H1 | SKLKGKSGPE |
| H1C11 | SKLKGKSGPEFHWFINVPPF |
| H1E2 | SKLKGKSGPEIRVLMSRVFA |

| | Mimotope sequence | $K_D$ (nM) |
|---|---|---|
| A2 | LLLSWDPWRIECLPP | 70.77 |
| A3 | TEAEGRLAWCRKWAYV | 23.01 |
| A6 | RMDSRFMVDWPCLNW | 79.85 |
| A11 | SYDFQCFPWCGH | 373.9 |
| C1 | GWNPWADIACSLP | 103.7 |
| C5 | ACSSFKTWPCWPE | 657.7 |
| E4 | SPWWDNDPPP | 22.48 |
| F5 | SSLSGPCKYCWYK | 440.7 |
| F6 | PRLSLCTIVLDCRGL | 95.96 |
| G6 | SMQLMRSDYGCPPR | 174.9 |
| H4 | YPKLRCVEWT | 293.6 |

**Extended Data Figure 4. Identification of mimotopes specific for anti-mouse CD19 and anti-ALK antibodies.** (**A-D**) Identification of a mimotope for anti-mouse CD19 mAb 1D3. (**A**) Flow cytometry scatter plots illustrating successful identification of yeast cells binding to $1D3^{IgG}$ at 1µM. Two yeast populations, P1 and P2, were identified. (**B**) Top-ranked 1D3 mimotopes. Shown in the table are individual mimotopes and mimotope fusions. (**C**) ELISA showing the binding of $1D3^{IgG}$ to chemically synthesized mimotopes in B. A one site-specific binding model was used for assessing the apparent binding affinity. (**D**) Mouse IFN-γ secretion from anti-mouse CD19 CAR-T cells co-cultured with target cells labeled with 100nM of amph-mimotopes or free mimotope peptides. Error bars show mean ± s.d. with three replicates. **, $p<0.01$ by one-way ANOVA with Turkey's post-test. (**E-F**) Identification of a mimotope for ALK-specific mAb ALK123. (**E**) Flow cytometry plots showing the successful identification of yeast cells binding to $ALK123^{IgG}$ at 500nM. (**F**) Binding of $ALK123^{IgG}$ to select yeast clones. Amino acid sequences were shown for each yeast clone. A one site-specific binding model was used for assessing the apparent binding affinity.

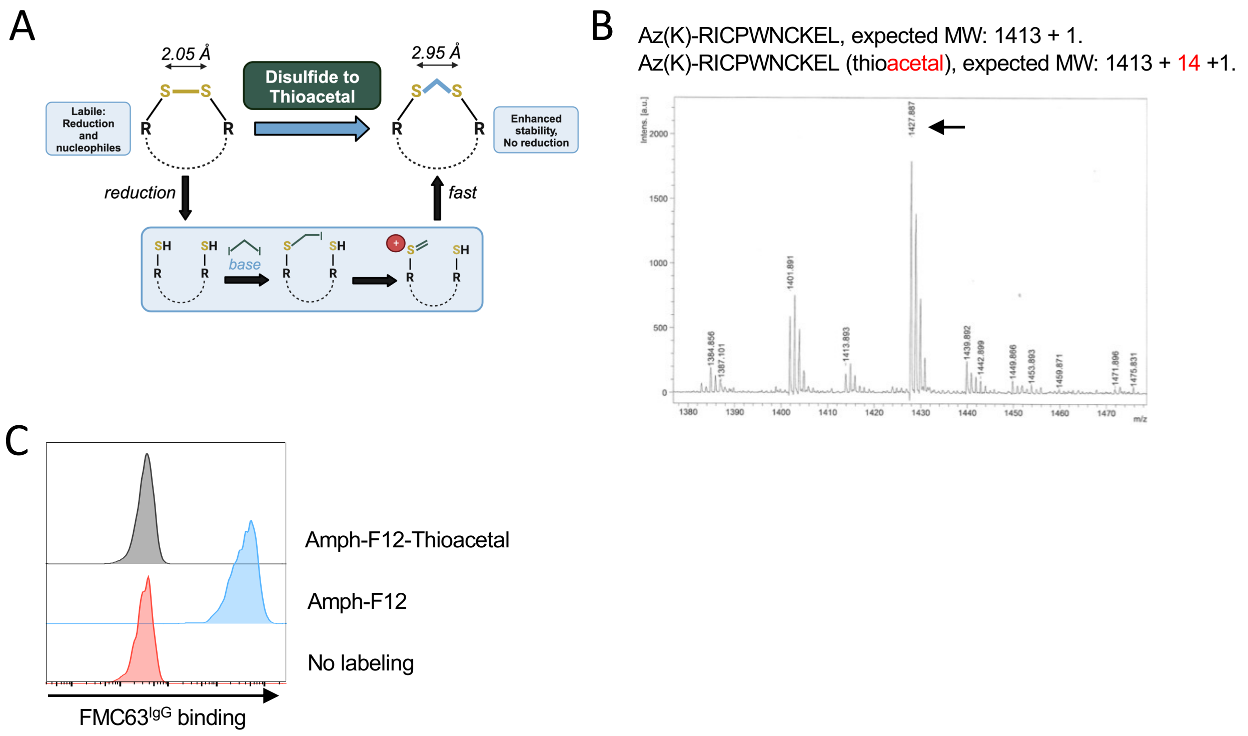


**Extended Data Figure 5. Converting the disulfide bond to a thioacetal bond prevents mimotope binding to FMC63$^{IgG}$.** (**A**) schematics showing the chemical structure and conversion of the disulfide bond to a thioacetal bond. (**B**) MALDI spectrum showing the purification of expected mimotope with a thioacetal bond. (**C**) Representative histogram showing FMC63$^{IgG}$ binding to target cells labeled with amph-mimotope$^{F12}$ variants.

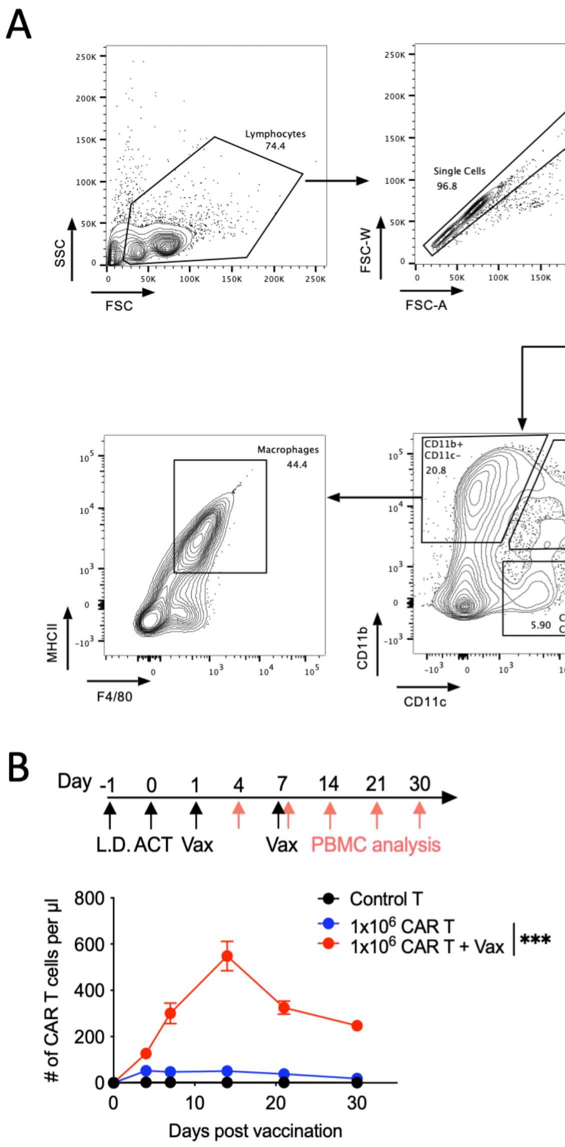


**Extended Data Figure 6. Amph-mimotope labeling of APCs and stimulation of FMC63-mCAR-T cell expansion *in vivo*.** (**A**) Gating strategies and surface makers used for defining APC populations in the lymph node. (**B**) C57BL/6 mice (*n*=5 animals/group) were lymphodepleted (L.D.) with 500cGy gamma irradiation, adoptively transferred with $10^6$ FMC63-mCAR-T cells, and then vaccinated at indicated time points. Shown are the number of circulating FMC63-mCAR-T cells per microliter of blood quantified by flow cytometry over time. This data is related to Fig. 5E. Error bars show mean ± 95% CI. ***, p<0.0001 by two-way ANOVA with Tukey's post-test.

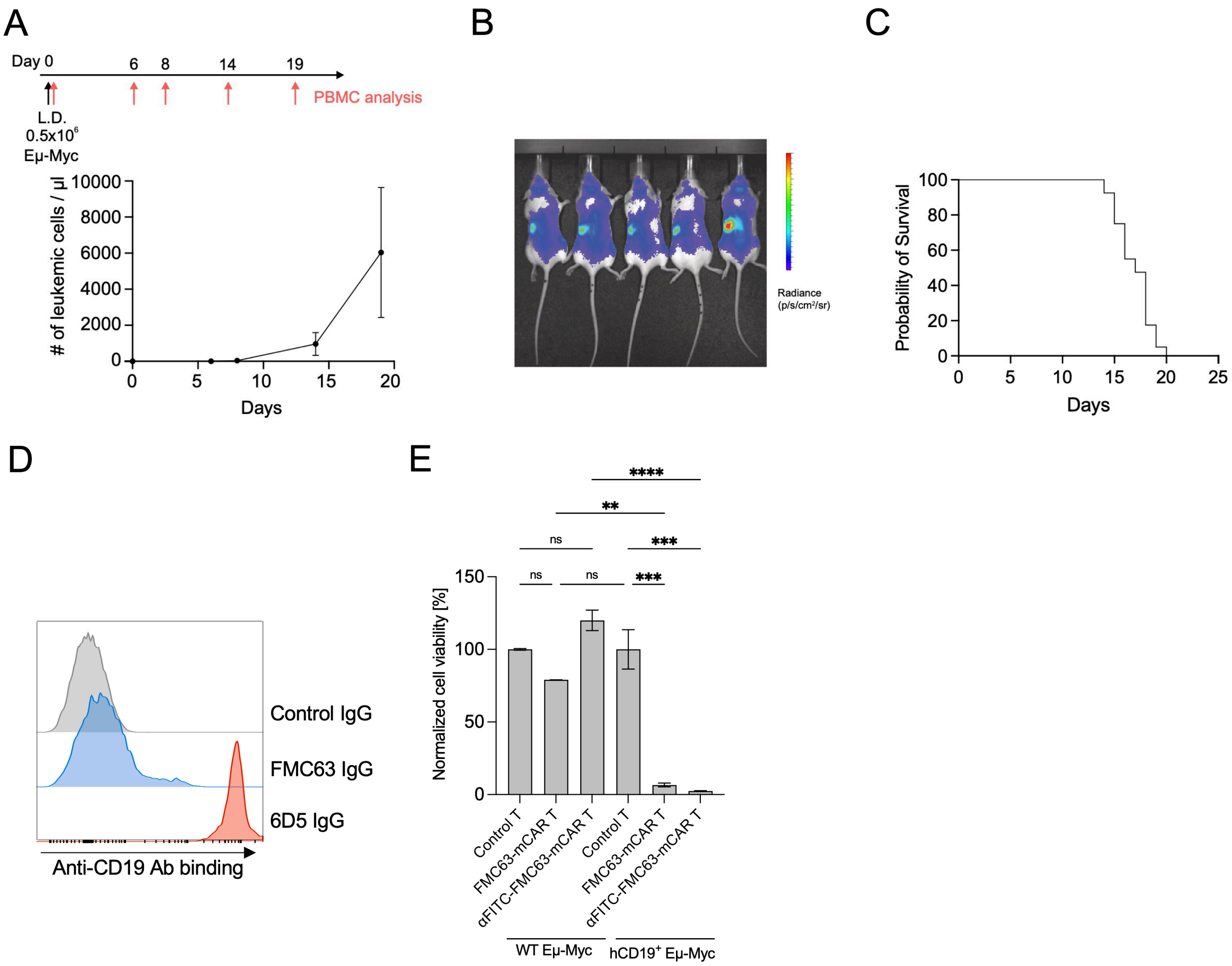


**Extended Data Figure 7. Establishment of the hCD19$^{+}$ Eμ-Myc B-ALL/Lymphoma mouse model for assessing hCD19-targeted mCAR-T therapy.** (**A**) Experimental setup and timeline and enumeration of circulating leukemic cells (n=4). C57BL/6 albino mice were lymphodepleted (L.D.) and injected with $0.5 \times 10^{6}$ hCD19$^{+}$ Eμ-Myc cells. Peripheral blood was collected on days 0, 6, 8, 14 and 19. (**B**) Representative IVIS imaging of Eμ-Myc albino mice at day 4. (**C**) Overall survival of untreated Eμ-Myc mice in this manuscript (n=40). (**D**) FMC63 IgG binding to primary mouse B cells as monitored using flow cytometry. An anti-mouse CD19 IgG clone 6D5 was included as a control. (**E**) Cytotoxicity of FMC63 mCAR-T or tandem αFITC-FMC63-mCAR-T against WT (murine CD19$^{+}$) or hCD19$^{+}$ Eμ-Myc mouse B-ALL cells. A luciferase-based assay was used for assessing cell killing by CAR-T (see Materials and Methods). Error bars show mean±95% CI for A, mean ± s.d. (n=2-3) for E. **, $p<0.01$; ***, $p<0.001$; ****, $p<0.0001$; ns, non-significant by one-way ANOVA with Tukey's post-test.

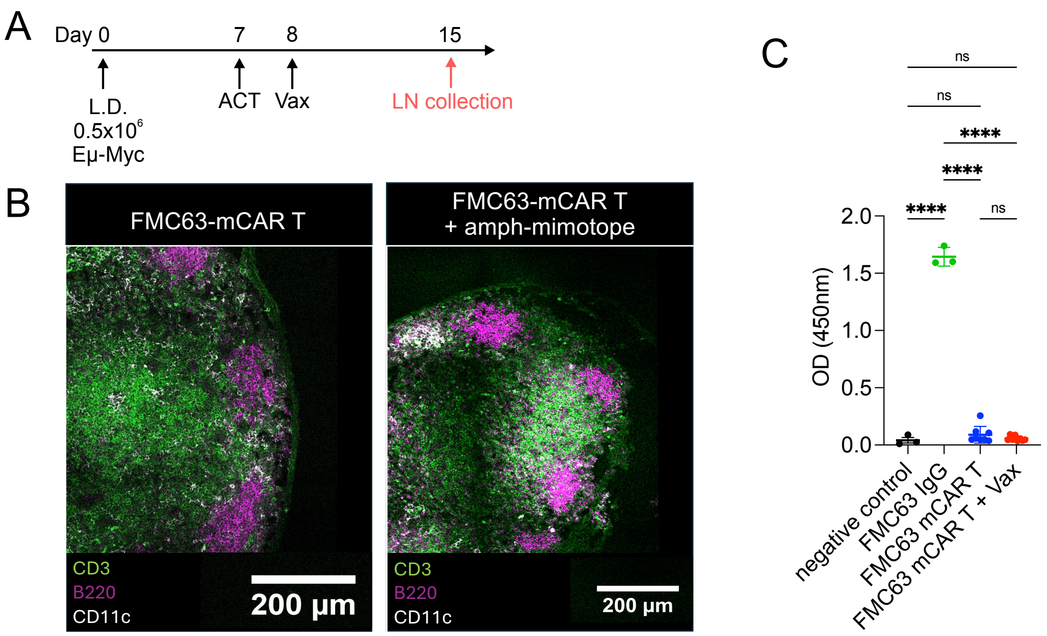


**Extended Data Figure 8. Evaluation of the impact of vaccine boosting of CAR-T cells on LN architecture and induction of serum antibodies against amph-mimotope vaccine.** (**A**) Experimental setup and timeline. C57BL/6 mice were lymphodepleted (L.D.) and injected with $0.5\times10^6$ $hCD19^+$ Eμ-Myc cells. On day 7, mice were adoptively transferred with $2\times10^6$ FMC63-mCAR-T cells or control T cells, then vaccinated 1 day later with 10μg amph-F12-A1 (Vax). Inguinal (draining) lymph nodes were collected on day 15, frozen, and stained for CD3, B220, and CD11c. (**B**) Confocal imaging of LNs. Shown are LN sections stained with anti-CD3, anti-B220, and anti-CD11c to define the T cell zone, B cell zone and DC populations in the LN. (**C**) Evaluation of serum antibody against amph-mimotope vaccine. Serum was collected from FMC63-mCAR-T- and FMC63-mCAR-T + Vax-treated Eμ-Myc-bearing mice on day 21 (timeline presented in **Fig. 6A**) for analysis of mimotope-specific IgG by ELISA (n=7 for FMC63-mCAR-T, n=8 for FMC63-mCAR-T + Vax). FMC63 IgG was used as a positive control. This data is related to Fig. 6. Error bars show mean±95% CI. ****, $p<0.0001$; ns, non-significant by one-way ANOVA with Tukey's post-test.

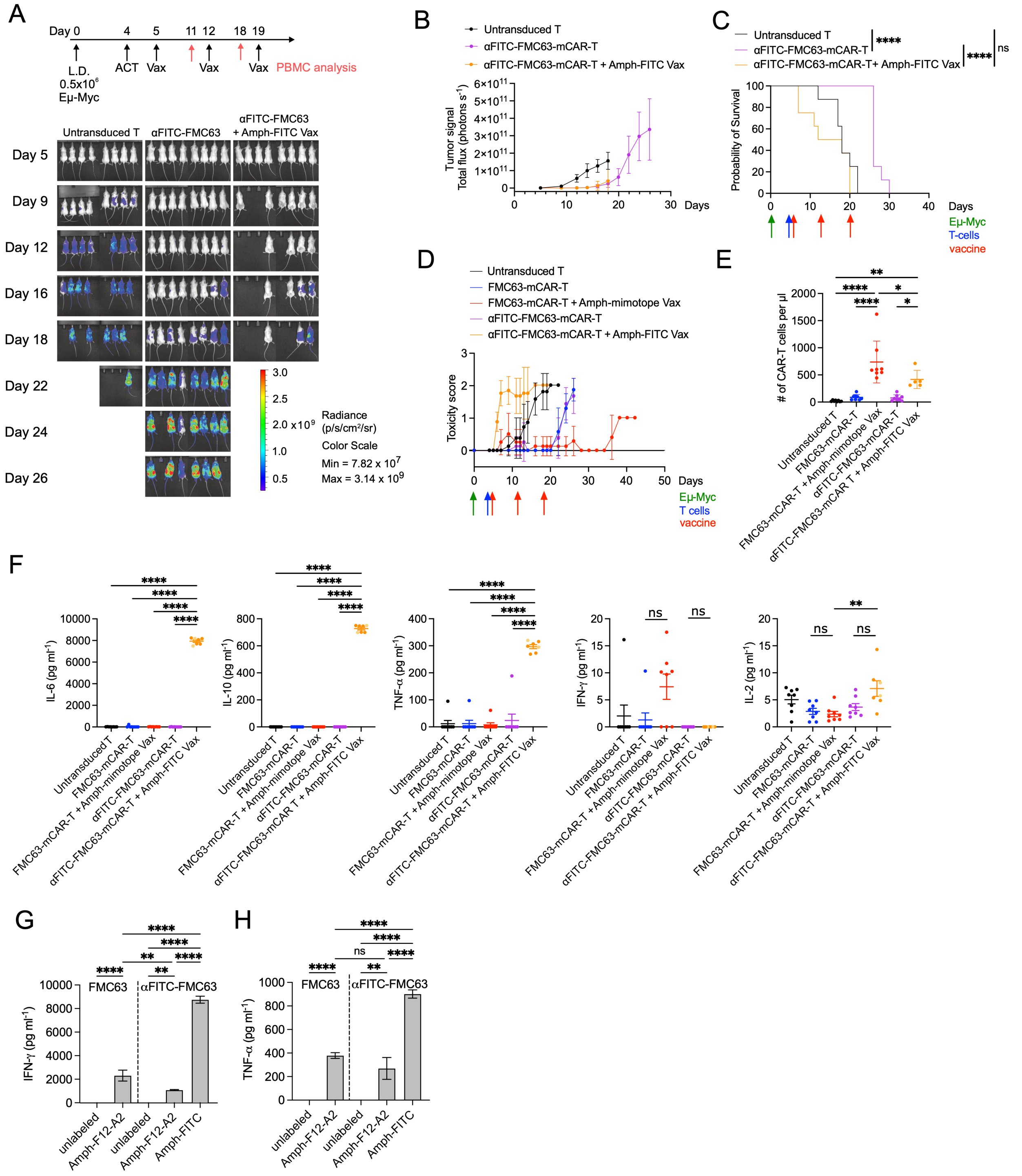


**Extended Data Figure 9. Toxicity of αFITC-FMC63-mCAR-T cells and amph-FITC vaccine therapy in a hCD19⁺ Eμ-Myc B-ALL/Lymphoma mouse model.** (**A**) Experimental setup and whole animal imaging of disease progression. B6 albino mice were lymphodepleted (L.D.) and injected with $0.5\times10^6$ hCD19⁺ Eμ-Myc cells. On day 4, mice were adoptively transferred with $2\times10^6$ αFITC-FMC63-mCAR-T cells or FMC63-mCAR-T (mCAR-T), then vaccinated 1 day later with 10nM amph-FITC (amph-FITC vax) or 10 μg amph-F12-A1 (amph-mimotope Vax) (n=8 animals/group). Vaccinations were repeated 7 and 14 days later. mFMC63-MCAR-T, and FMC63-

mCAR-T + Amph-mimotope Vax are the same as presented in Fig. 6. (**B**) Quantification of total photon counts over time in different treatment groups. (**C**) Eμ-Myc-injected mice were monitored for overall survival. (**D**) Toxicity score of each treatment group. Mice were monitored daily for clinical signs of toxicity before CAR-T injection and until mice reached the endpoint of the study. The toxicity of therapy was monitored using a toxicity score as described before[21]. A toxicity score of 0 corresponded to active, well-groomed animals, with well-kept hair coat and invisible spine. A score of 1 was assigned to mice with signs of hypomotility, tousled hair coat and a partially visible spine. A score of 2 was given in case of somnolence, rough, dull or soiled hair coat, hunched back with visible spine. (**E**) Enumeration of circulating CAR-T cells or control $CD45.1^+$ T cells by flow cytometry on day 11. Control mice, mFMC63-MCAR-T, and FMC63-mCAR-T + Amph-mimotope Vax are the same as presented in **Fig. 6C**. (**F**) Serum cytokine analysis. Blood was collected on day 11 (solid yellow dots) or day 7 for amph-FITC-treated mice (light yellow dots), and ELISA was performed to determine the concentration of IL-6, IL-10, TNF-α, IFN-γ, and IL-2 in the serum. (**G-H**) Cytokine responses of CAR-T cells to amph-vax labeled K562 cells. FMC63-mCAR-T and αFITC-FMC63-mCAR-T were co-cultured with K562 labeled with 500nM Amph-F12-A2 or 500nM Amph-FITC at a 10:1 E:T ratio for 6 hours followed by a measurement of IFN-γ and TNF-α concentration in supernatant by ELISA. Error bars show mean±95% CI for C, E, F; or mean±s.e.m. for G, H. *, $p<0.05$, **, $p<0.01$; ***, $p<0.001$; ****, $p<0.0001$; ns, non-significant by Long-rank (Mantel-Cox) test for C, one-way ANOVA with Tukey's post-test for E, F, G by two-way ANOVA with Tukey's post-test for G, H.

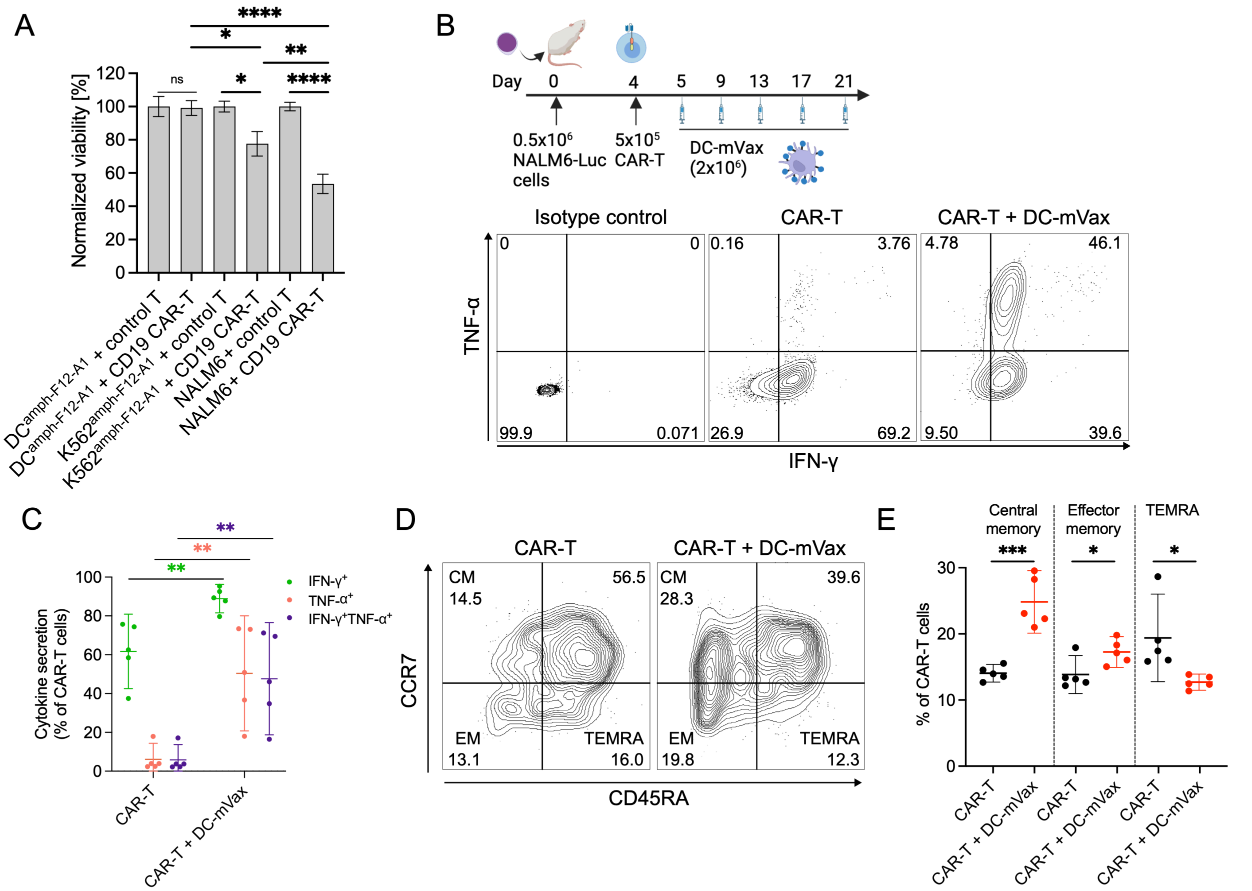


**Extended Data Figure 10. Resistance of amph-F12-A1-labeled DC to killing by CAR-T cells and immunophenotype of CAR-T cells boosted with DC-mVax.** (**A**) Sensitivity of DC-mVax to CAR-T cells. Activated MoDCs or K562 cells were incubated with 500 nM amph-F12-A1 for 30 min, washed, and then co-cultured with human CD19 CAR-T cells or control untransduced T cells at a 10:1 E:T ratio for 6 hours followed by measurement of target cells viability by flow cytometry (n=5). (**B-C**) CAR-T cell cytokine polyfunctionality analysis. 0.5 x $10^6$ CAR-T cells were administered with or without DC-mVax as shown in the timeline (n=5 animals/group). Spleens were harvested on day 19. Splenocytes were incubated for 6 hours with a cell stimulation cocktail followed by intracellular staining with anti-IFN-γ and anti-TNF-α antibodies. Shown are the representative contour plots (**B**) and the percentage of IFN-$\gamma^+$, TNF-$\alpha^{+}$, and IFN-$\gamma^+$TNF-$\alpha^+$ CAR-T population (**C**). (**D-E**) Immunophenotyping of CAR-T cells in the spleen on day 19. Shown are representative contour plots (**D**) and the percentage of CAR-T (**E**) with CD45RA$^-$CCR7$^+$ central memory (CM), CD45RA$^-$CCR7$^-$ effector memory (EM), and CD45RA$^+$CCR7$^-$ effector memory re-expressing CD45RA (TEMRA) T cells phenotype. Error bars show mean ± s.e.m. for A, or mean ± 95% CI. for C, E. *, p<0.05; **, p<0.01; ***, p<0.001; ****, p<0.0001; ns, non-significant by two-way ANOVA with Tukey's post-test for A, by unpaired t-test for C, E.
